# Neuronal overexpression of potassium channel subunit Kcnn1 prolongs survival of SOD1-linked ALS and A53T alpha-synuclein mouse models

**DOI:** 10.1101/2024.10.11.617887

**Authors:** Maria Nagy, Justin Cotney, Wayne A. Fenton, Arthur L. Horwich

## Abstract

Eye muscles and the motor neurons in the innervating cranial nerve nuclei are relatively spared in human ALS, and likewise, these cranial motor neurons are spared of SOD1YFP aggregation in a transgenic mouse model of SOD1-linked ALS, G85R SOD1YFP. RNA profiling of mouse oculomotor (CN3) neurons (resistant) vs hypoglossal (CN12) and spinal cord motor neurons (susceptible) from nontransgenic mice identified differentially expressed channel and receptor genes. A number were evaluated for effects on survival of the ALS strain by transgenesis or knockout to emulate the relative RNA level in oculomotor neurons. Transgenesis of Thy1.2-driven cDNA for mouse Kcnn1, a potassium channel subunit, extended the median days of survival time to paralysis of mutant G85R SOD1YFP mice by up to 100%, associated with absence of fluorescent aggregates; extended the median time to paralysis of G93A SOD1 mice by up to 55%; and extended the median time to endstage motor disease of a Thy1.2-driven alpha-synuclein transgenic strain by up to greater than 100%. The overexpressed Kcnn1 subunit was diffusely cytoplasmic in motor neurons and found to induce a multifaceted stress response as judged by RNAseq and immunostaining, including ER stress response, mitochondrial stress response, and an integrated stress response. Like other potassium channel subunits, Kcnn1 subunit is likely targeted to the ER, but as reported earlier in rodent Kcnn1-transfected cultured cells, in the absence of Kcnn2 with which to co-assemble, Kcnn1 is channel-inactive and is diffusely cytoplasmic. Thus, a nonassembled and potentially misfolded state of overexpressed Kcnn1 targeted to the ER of neurons may explain the stress responses, which in the mutant SOD1 and A53T alpha-synuclein mice, protect against the pathogenic proteins.

Major neurodegenerative diseases, including Alzheimer’s Disease and Parkinson’s Disease, are associated with the accumulation of characteristic proteins, Abeta/Tau and alpha-synuclein in AD and PD, respectively, that misfold, aggregate, and in many cases form amyloid fibrils (e.g. Long and Holtzman, 2019; Sierksma et al, 2020; Tanner et al, 2024). Such pathogenic behavior is associated with malfunction/death of specific neuronal populations, producing consequent clinical symptoms. It seems counterintuitive to observe proteinopathy as a major facet of these diseases considering that there is generally a quality control machinery in all cells, consisting of effectors - molecular chaperones, ubiquitin/proteasomal components, and autophagy/lysosome components - governed by a “sensor” circuitry – e.g. UPR, ISR, HSF - that can detect such misbehavior and induce protective responses. While neurons may be particularly susceptible because they are postmitotic and unable to distribute damaging protein species to daughter cells as a protective means, it has remained unclear whether the endogenous sensor/effector pathways can be induced sufficiently in vivo so as to mediate protection. Here, we report that neuronal overexpression of a potassium channel subunit, mouse Kcnn1, in two different transgenic mouse neurodegenerative models, protects against aggregation and cell loss by apparent induction of multiple stress response pathways, substantially extending survival of the mice.

## RESULTS

### Transgenic Thy1.2-driven Kcnn1 extends survival time to paralysis of G85R SOD1YFP transgenic mice

In earlier studies of transgenic G85R mutant SOD1YFP mice, a model for ALS, we observed paralysis by 6-7 months of age, associated with large aggregates of SOD1YFP appearing in spinal cord motor neurons by 2-3 months of age (Hadzipasic 2014). Consonant with clinical observations of sparing of extraocular muscles and their innvervating cranial motor nuclei in both mutation-associated and sporadic human ALS patients, we observed sparing of aggregation in motor neurons innervating the extraocular muscles of G85R SOD1YFP mice, CN3 (oculomotor), CN4 (trochlear), and CN6 (abducens) nuclei, termed “resistant”, while large aggregates were observed in motor neurons of CN7 (facial) and CN12 (hypoglossal) nuclei, as well as in motor neurons in spinal cord ventral horn, termed “susceptible” (Thomas et al., 2018). We carried out laser capture microdissection of motor neurons from CN3, CN12, and spinal cord (SC) of a normal B6SJL mouse, recovered total RNA, and carried out RNAseq. Differential expression analysis revealed that amounts of transcripts for a number of channel and neurotransmitter receptor proteins were prominently different between CN3 (resistant) and CN12/SC (susceptible). We hypothesized that reducing these differences for individual transcripts might convert susceptible neurons to resistant ones and, hence, positively affect survival of both neurons and animal. In particular, where RNA abundance was less in resistant CN3 than in susceptible SC/CN12, we produced mouse strains with knockouts of these genes by CRISPR/Cas techniques; and where transcript abundance was greater in resistant CN3 than in susceptible SC/CN12, we produced transgene strains with Thy1.2 promoter-driven mouse cDNAs (directing neuronal expression, Caroni, 1997). These were all crossed into the G85R SOD1YFP homozygous line and survival time to paralysis determined. None of 6 selected knockouts nor 5 of 6 transgenics showed any prolongation of survival relative to G85R SOD1YFP (see Kaplan-Meier plots in SF1, SF2). However, a transgenic strain carrying heterozygous Thy1.2-driven mouse Kcnn1 (copy number ranging from 3-4 copies by real-time PCR) showed a significant increase in survival of G85R SOD1YFP homozygous mice (with copy numbers between 260 and 330; Fig.1A, red trace). The median survival was 267 days, increased from 207 days for control G85R SOD1YFP homozygote littermates (blue trace), amounting to 29% extension of survival (p = 10^-6^), and the longest survival was 330 days. Littermates lacking this so-called Kcnn1-3 transgene exhibited a survival that closely resembled that of the cohort of G85R SOD1YFP homozygous mice (compare blue with black traces in Fig.1A).

**Figure 1.**
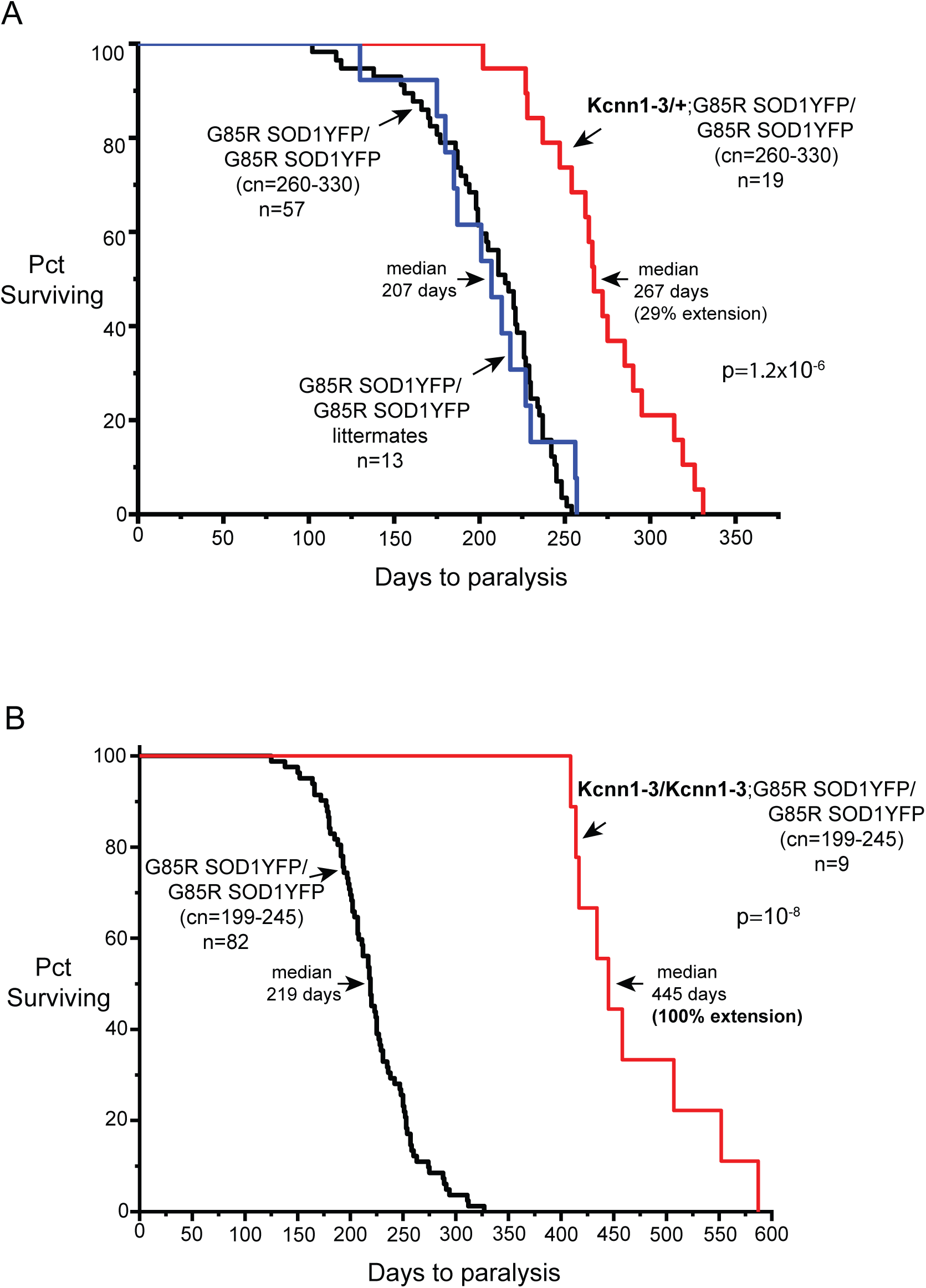
Kaplan-Meier survival plots for G85R SOD1YFP homozygous mice of high copy number, 260-330, (panel A), and lower copy number, 199-245, (panel B), showing that survival (days to paralysis) is extended by Thy1.2-Kcnn1 transgenesis by 35% and 100%, respectively. Extended survival of Kcnn1 transgenic mice (red trace) is shown relative to both littermates lacking Kcnn1 (blue trace in A) and a cohort of homozygous G85R SOD1YFP mice (black trace). Note that Thy1.2-Kcnn1 is hemizygous in (A) and homozygous in (B), but that, in the case of G85R SOD1YFP, there was no significant difference in survival for copy number 260-330 between hemizygous and homozygous Kcnn1 (compare panel (A) with SF3).

We tested whether homozygosis of Kcnn1-3 would further improve survival and observed only a very small increase of median survival (65 days extension vs 60 days; SF3). Because the production of mutant SOD1-linked neurodegeneration in mice typically requires high transgenic copy number, whereas humans are typically affected by a single copy, we asked whether survival would be further extended if we were to maintain the Kcnn1 copy number (at 6-8, via homozygosity) while lowering the G85R SOD1YFP copy range from 260-330 down to 199-245. We observed that median survival of the control mice remained about the same, at 219 days, but that now the median survival in the presence of homozygous Kcnn1-3 was doubled to ∼445 days (100% extension), with longest survival now correspondingly extended to nearly 600 days (Fig.1B). This implies that Kcnn1 transgenesis has more profound effects when the SOD1 copy number is lower and raises the question of what copy level of Kcnn1 would be sufficient to protect against a single copy of mutant SOD1 in the human setting.

### Transgenic Thy1.2-Kcnn1 also extends survival time of G93A mice

We next tested a more severe allele of SOD1, G93A (Gurney et al, 1994), which at hemizygosity, with copy number ranging from ∼180-220, produces paralysis in a B6SJL background by ∼120 days. Here, we observed only modest effects of Kcnn1-3 transgenesis, extending survival by a median of ∼10-15 days. However, when we homozygosed Kcnn1-3 (achieving a Kcnn1 copy number of 7-8), a striking extension of G93A survival was observed, with a median of 190 days compared with 124 days in the absence of Kcnn1, amounting to 54% extension of survival and with the longest surviving mouse extending by nearly 100 days (Fig.2A). To exclude that this was a potential insertional effect, an independent Thy1.2-Kcnn1 transgenic line was produced (called Kcnn1-6), with a hemizygous copy number of 6. When crossed with G93A, these mice also exhibited an extended survival, in this case with a median of 166 days, as compared with 125 days in the absence of Kcnn1, an extension of 34% (Fig.2B). Here, the longest-surviving mouse was found to reach ∼225 days, consistent with an outlier Kcnn1 copy number of 10. Thus, it appears that the effect of Kcnn1 transgenesis is independent of the insertion site and that increased copy number of Kcnn1 has a larger effect on survival of G93A mice.

**Figure 2.**
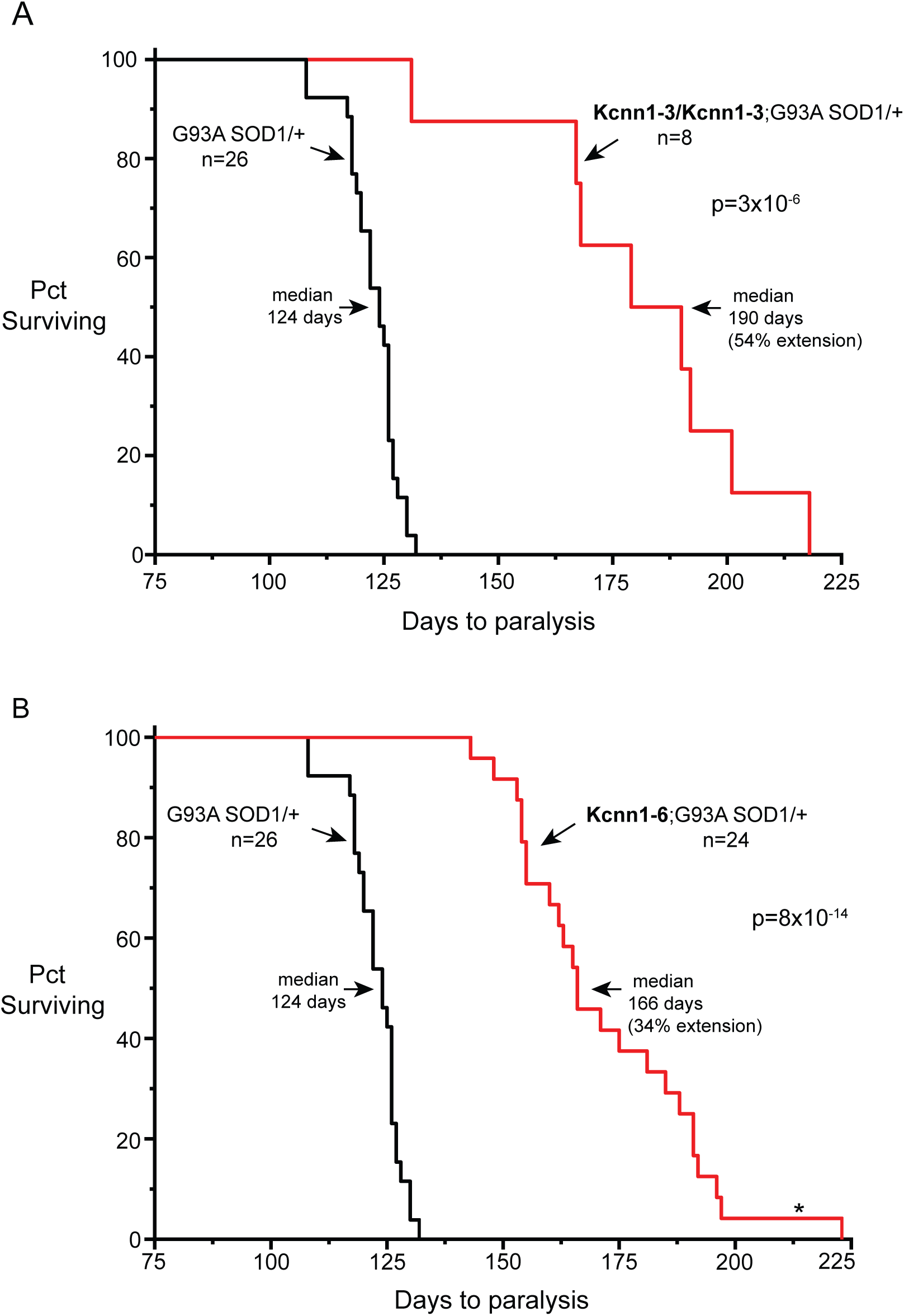
Kaplan-Meier survival plots for G93A SOD1/+ mice (cohort of 26 mice in both panels A and B) and G93A SOD1/+ mice homozygous for Kcnn 1-3 (totaling ∼8 copies by real time PCR measurement) and a separate hemizygous Kcnn1-6 line (with 6 copies by real time PCR) (red traces). Extension of time to paralysis was 54% in panel A and 34% in panel B.

Notably, there are 4 homologous Kcnn genes in mouse and man (Kohler et al, 1996; Adelman et al, 2012). Subunits expressed from these genes assemble into tetrameric calcium-activated potassium channels that have calmodulin bound constitutively at the C-termini of their subunits. They function at the afterhyperpolarization phase of action potentials (Yost, 1999), with calcium entering via N-type channels mediating an allosteric opening of the central K^+^ channel (see Lee and MacKinnon, 2018 for structural analysis/mechanism of activation). Kcnn1, Kcnn2, and Kcnn3 gene products form small conductance (SK) channels while Kcnn4 forms an intermediate conductance (IK) channel. Because Kcnn2 has nearly 90% predicted amino acid identity to Kcnn1 through its channel and calmodulin binding regions and is coexpressed with Kcnn1 in cortex and hippocampus (Stocker, 2004), we asked whether a transgene with Thy1.2-driven mouse Kcnn2 cDNA could produce the same improved survival effects in G85R SOD1YFP homozygous mice as Kcnn1. We produced four Thy1.2-Kcnn2 transgenic lines, but observed that none of them exhibited any improvement of survival of the G85R SOD1YFP homozygous strain, despite obtaining copy numbers for two lines that were far greater than those achieved with Thy1.2-Kcnn1 mice (SF4). Thus the beneficial effect observed here of Thy1.2-mouse Kcnn1 transgenesis is not shared by Kcnn2.

### Increased expression of Kcnn1 transcript and encoded subunit in spinal cord motor neurons of Kcnn1 transgenic mice

To assess whether the survival effects of Kcnn1 transgenesis were mediated at the level of motor neuron expression, spinal cord sections of a homozygous Kcnn1-3 mouse were assessed first by in situ hybridization with a Kcnn1 RNA probe. This detected cytoplasmic signals in the soma of large motor neurons in the ventral horn that were ∼5- 10-fold greater in intensity than in control B6SJL spinal cord (Fig.3A). Next, a polyclonal antibody specific to the encoded Kcnn1 subunit (against the 150 aa C-terminal portion) was used to probe spinal cord sections. Signals were observed within the cytoplasm of large ventral horn motor neurons, as well as from a few smaller cells, with an ∼5-10 fold elevation of intensity relative to nontransgenic spinal cord (Fig.3B). The signal seemed to be excluded from the nucleus (or, at minimum, from the nucleolus) and extended into neuronal processes. Antibody was also used to probe Kcnn1-6 spinal cord and here also produced an intracellular staining pattern, with clearcut nuclear exclusion (Fig.3C). Immunostaining intensity relative to nontransgenic was once again ∼5-10-fold greater. We also inspected motor cortex by immunostaining and observed a similar staining pattern for Kcnn1 product in layer V motor neuron soma (SF5). Thus, it appears that in transgenic Thy1.2-Kcnn1 mice, the Kcnn1 cDNA is overexpressed at both mRNA and protein levels in upper and lower motor neurons, supporting that they are likely the major site of benefit to survival of the mutant SOD1 mice.

**Figure 3.**
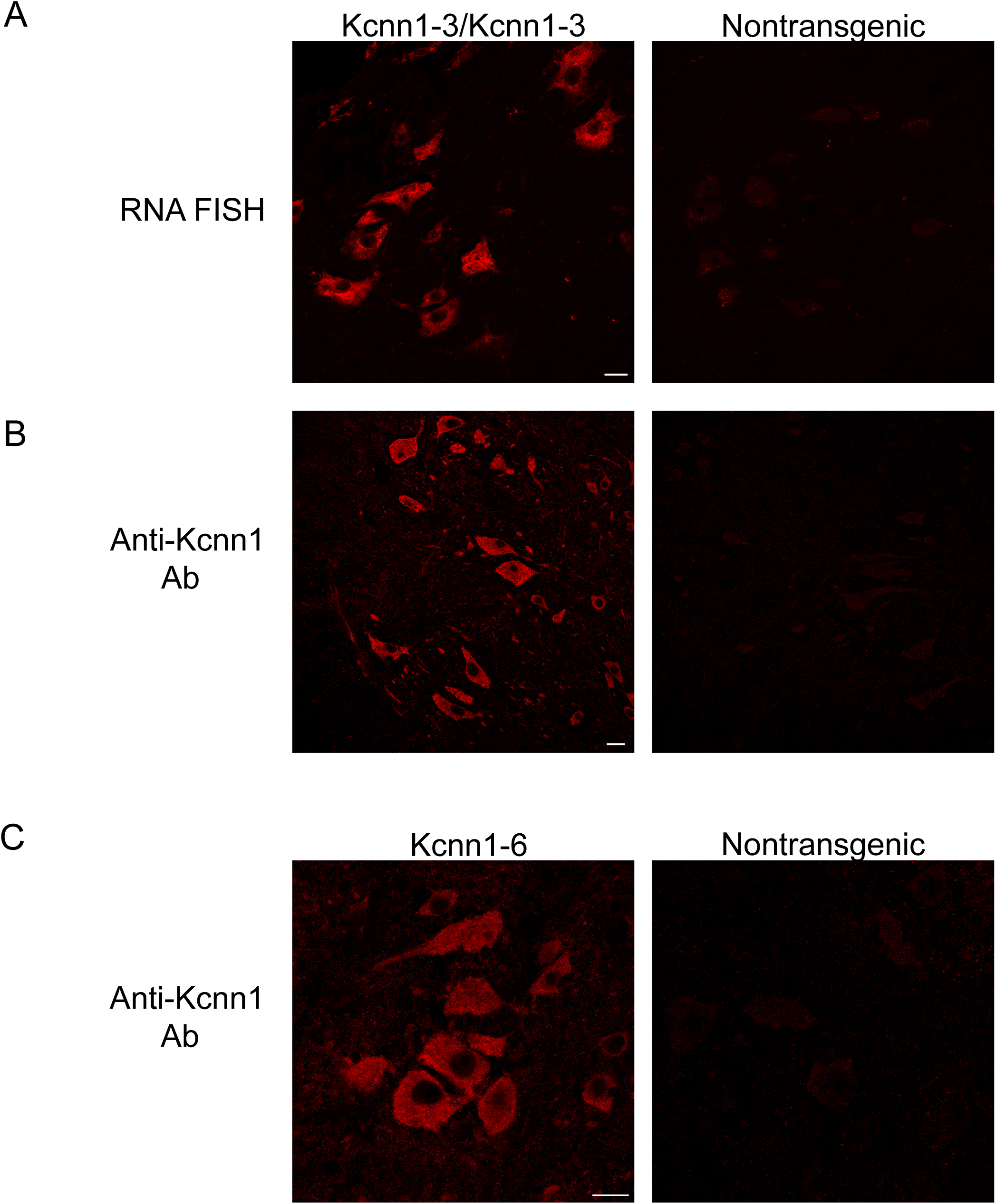
Expression of Kcnn1 RNA and protein in motor neurons in ventral horn region of Kcnn1-3/Kcnn1-3 homozygous mice and nontransgenic B6SJL. A, RNA fluorescence in situ hybridization (FISH) of Kcnn1 RNA present in ventral horn of cervical spinal cord section of a 7 month old Kcnn1-3/Kcnn1-3 mouse (left) and nontransgenic B6SJL mouse (right). B, Immunostaining with an anti-Kcnn1 antibody of the Kcnn1-3/Kcnn1-3 (left) and B6SJL (right). C, Immunostaining of a Kcnn1-6 mouse (left) and a B6SJL nontransgenic mouse (right). Scale bars are 20 μm.

Doubly homozygous Kcnn1-3/Kcnn1-3;G85R SOD1YFP/G85R SOD1YFP mice were also immunostained with an anti-Kcnn1 antibody, to inspect for the effect of transgenic Kcnn1 on mutant SOD1YFP. A typical ventral horn motor neuron is shown in Fig.4. Anti-Kcnn1 staining (top panel) was cytoplasmic in location and patchy in character, with areas of increased and decreased intensity. No clear rim of cell surface staining could be observed. YFP fluorescence (middle panel) corresponding to G85R SOD1YFP protein was more uniform in the cytoplasm, with a few patchy areas of decreased intensity. There was a perinuclear “rim” of YFP fluorescence. [Note that the G85R mutant form of SOD1YFP is generally excluded from the nucleus, whereas wt SOD1YFP is imported and observable in motor neuron nuclei (Wang et al, 2009)]. The merged image (bottom panel) suggested considerable regional overlap of the staining patterns, without precise colocalization (i.e. no yellow), but other areas showed only YFP fluorescence. Efforts to demonstrate overlap of Kcnn1 staining with specific organelle markers were not successful, nor could we establish whether Kcnn1 staining was specifically associated with membrane structures within the cytosolic compartment. When an axonal process that had been partly longitudinally transected was examined (SF6), we observed signals from the cut surface, with fairly uniform G85R SOD1YFP fluorescence but punctate anti-Kcnn1 staining. An axonal cross-section (SF6) showed YFP fluorescence both within the axoplasm and at the perimeter, while Kcnn1 subunit staining was observed only at the perimeter in a punctate pattern. The 6 TMs in the Kcnn1-encoded subunit may favor, whether assembled or not, lodging with lipid-rich structures such as axolemma or myelin sheath. Finally, Kcnn1 immunostaining was observed in the presynaptic region of neuromuscular junctions from lumbrical muscles, identified via post-synaptic staining with bungarotoxin (SF6). Thus the beneficial effects of Kcnn1 overexpression could be exerted at the level of soma, neuronal processes, or synapses.

**Figure 4.**
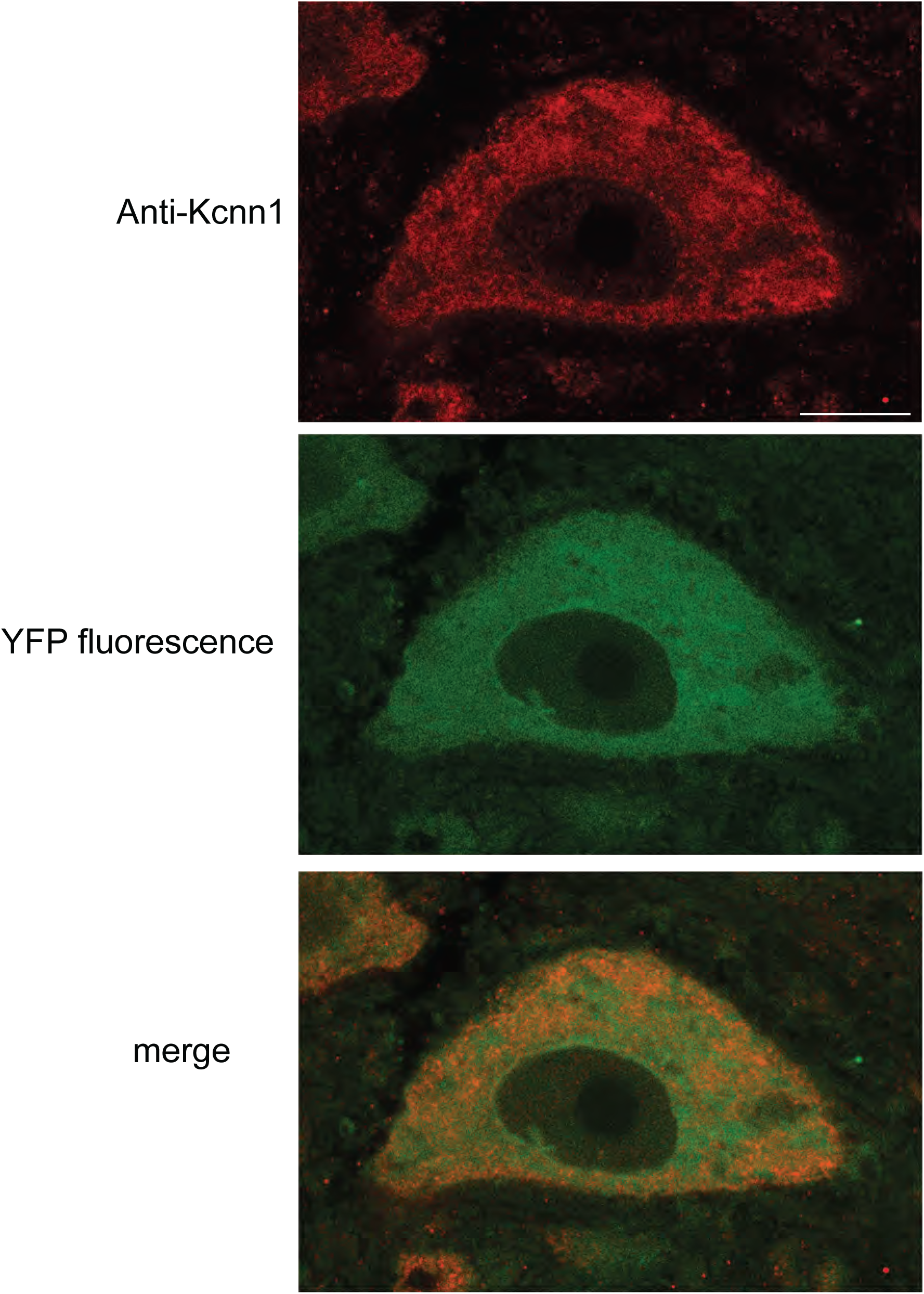
Comparison in a ventral horn neuron of Kcnn1 antibody staining pattern with G85R SOD1YFP homozygote fluorescence pattern. A 20 micron thick cervical spinal cord section from a 3 month old Kcnn1-3/ Kcnn1-3; G85R SOD1YFP homozygous mouse with copy number 298 was stained with anti-Kcnn1 antibody (top panel); the YFP fluorescence imaged (middle panel); and the two images merged (bottom panel). The Kcnn1 pattern was cytoplasmic and patchy, with variable intensity. The YFP fluorescence was more uniform with a rim of nuclear staining (see text), and the merge showed overlap of the patterns, but no distinct yellow color that would reflect precise colocalization. Scale bar is 10 µm.

Notably absent during examination of spinal cord sections of several double homozygous mice of 2-4 months of age (i.e. Kcnn1-3/Kcnn1-3; G85R SOD1YFP/G85R SOD1YFP) were lake-like aggregates (bright YFP-fluorescent, 30-50 Angstrom diameter structures within the soma; see e.g. Hadzipasic et al, 2014) that were invariably observed in a fraction of the motor neurons of homozygous G85R SOD1YFP mice of corresponding age. Thus, overexpression of Kcnn1 appears to prevent such aggregation.

Overexpression of Kcnn1 likewise prevented aggregation in the non-eye cranial motor nerve nuclei, facial (CN7) and hypoglossal (CN12). In the absence of the Kcnn1 transgene (Fig.5, top panels under 7N and 12N), florid aggregation was observed in the motor neurons in these nuclei in G85R SOD1YFP homozygous mice at 3 months of age (see also Thomas et al, 2018). In the presence of Kcnn1-3/Kcnn1-3, however, with notable overexpression of Kcnn1 observed by immunostaining (Fig.5, right panels), no aggregates were observed (lower panels under 7N and 12N; see also inset views).

**Figure 5.**
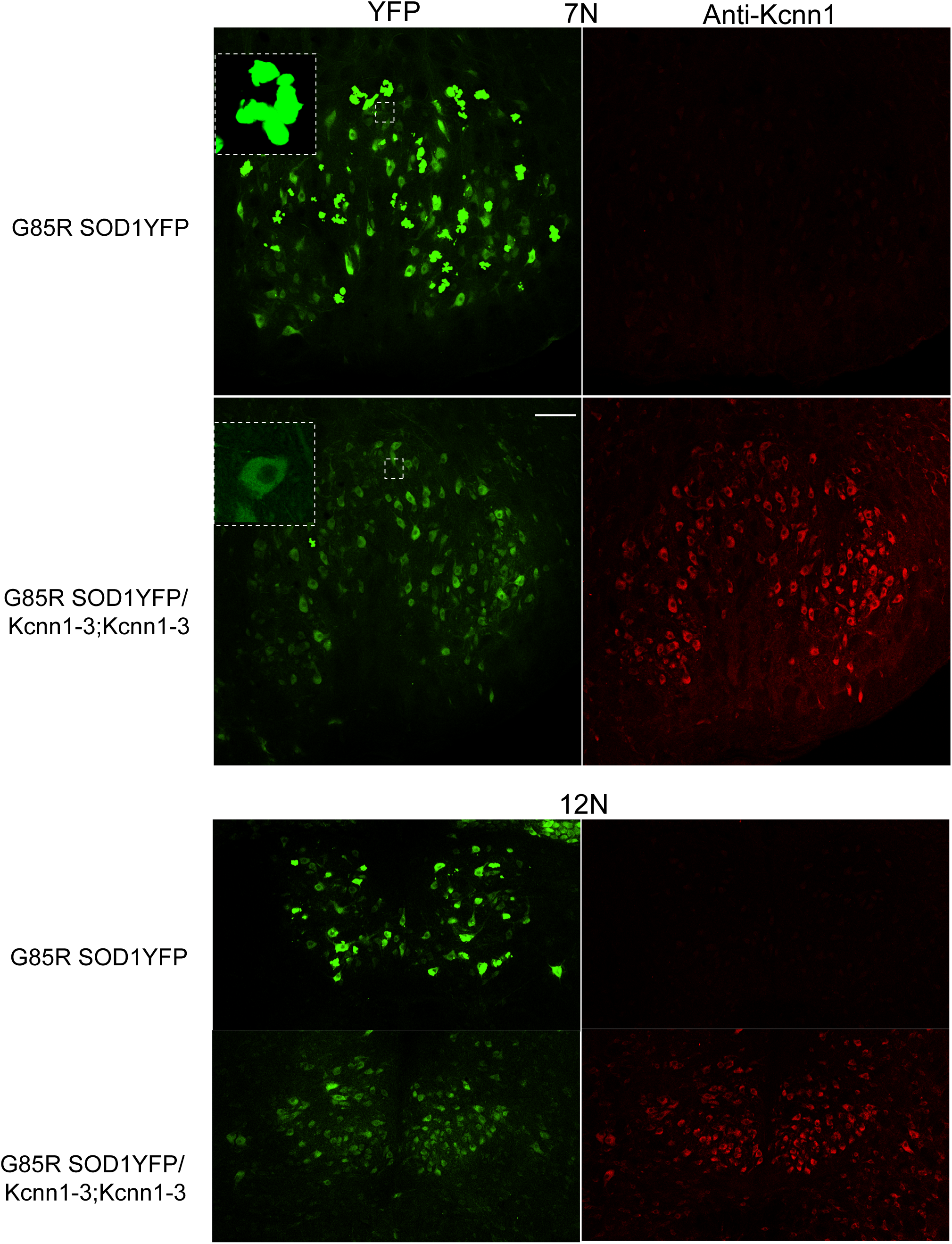
Non-ocular cranial motor neurons innervating facial muscles (7N) and tongue (12N) are prevented from accretion of SOD1 YFP aggregates by overexpression of Kcnn1 in double homozygote transgenics G85R SOD1YFP/G85R SOD1YFP;Kcnn1-3/Kcnn1-3. Left panels, YFP fluorescence from G85R SOD1YFP protein; right panels, immunostaining with anti-Kcnn1 antibody. In the absence of Kcnn1, YFP fluorescent aggregates are present in as many as 50% of motor neurons in 7N (top left panel, insert), and in an even higher percentage in 12N, but in the presence of overexpressed Kcnn1, detected in the right panels where the Kcnn1-3 homozygous transgene is present, only a rare YFP fluorescent aggregate (or artifact) could be observed.

### Intrathecal injection of newborn G93A pups with AAV9-CMV-Kcnn1 extends survival

As a further proof of principle, we assessed whether rAAV9-mediated delivery of Kcnn1 cDNA could affect the survival time to paralysis of G93A mice. In an initial test, we injected both lateral ventricles of P0 B6SJL mouse pups with a recombinant AAV9 encoding a CFP fluorescent marker, self-complementary AAV9-CMV-CFP (at 1.2 × 10E13 vp/ml), and observed 2-6 months later that there was substantial CFP fluorescence in ∼50-70% of ChAT-positive ventral horn motor neurons (SF7). A similar self-complementary AAV9 virus with CMV driving Kcnn1 was prepared at a higher titer, 4.5 × 10E13 vp/ml, and similarly injected into P0 pups from a mating of ∼2-3 month old G93A/B6SJL males with B6SJL females. The injected mice were subsequently genotyped, and 8 progeny that were G93A positive were followed to the point of paralysis. The Kaplan-Meier survival curve of these mice was compared with that of a cohort of uninjected G93A mice (Fig.6). This revealed non-overlapping survival curves, with the median survival of the injected mice extended by 19 days, a 15% extension of survival (p= 4.4X10^-6^). Fresh frozen spinal cord tissue pieces of ∼3 mm length were obtained from the injected G93A mice at the time of paralysis, and analyzed by qRT-PCR for the amount of Kcnn1 RNA relative to GAPDH RNA; this ratio was compared with that from a non-injected B6SJL control. We observed a direct correlation of survival time of the virus-injected G93A mice with the fold-increase of Kcnn1 RNA (SF8). Note that the highest level observed, ∼140-fold, approaches the levels observed with transgenic Kcnn1-6 (∼200-fold), and that survival time, ∼155 days, approaches the median survival time of Kcnn1-6/+;G93A/+ constitutively transgenic mice (166 days, Fig.2B). Spinal cord cross-sections of the mouse surviving 155 days were immunostained with anti-Kcnn1 antibody, confirming overexpression of the protein in motor neurons relative to those of an uninjected B6SJL mouse (SF9), revealing also that virtually 100% motor neuron transduction of ChAT positive large neurons had occurred in vental horn following the P0 injection.

**Figure 6.**
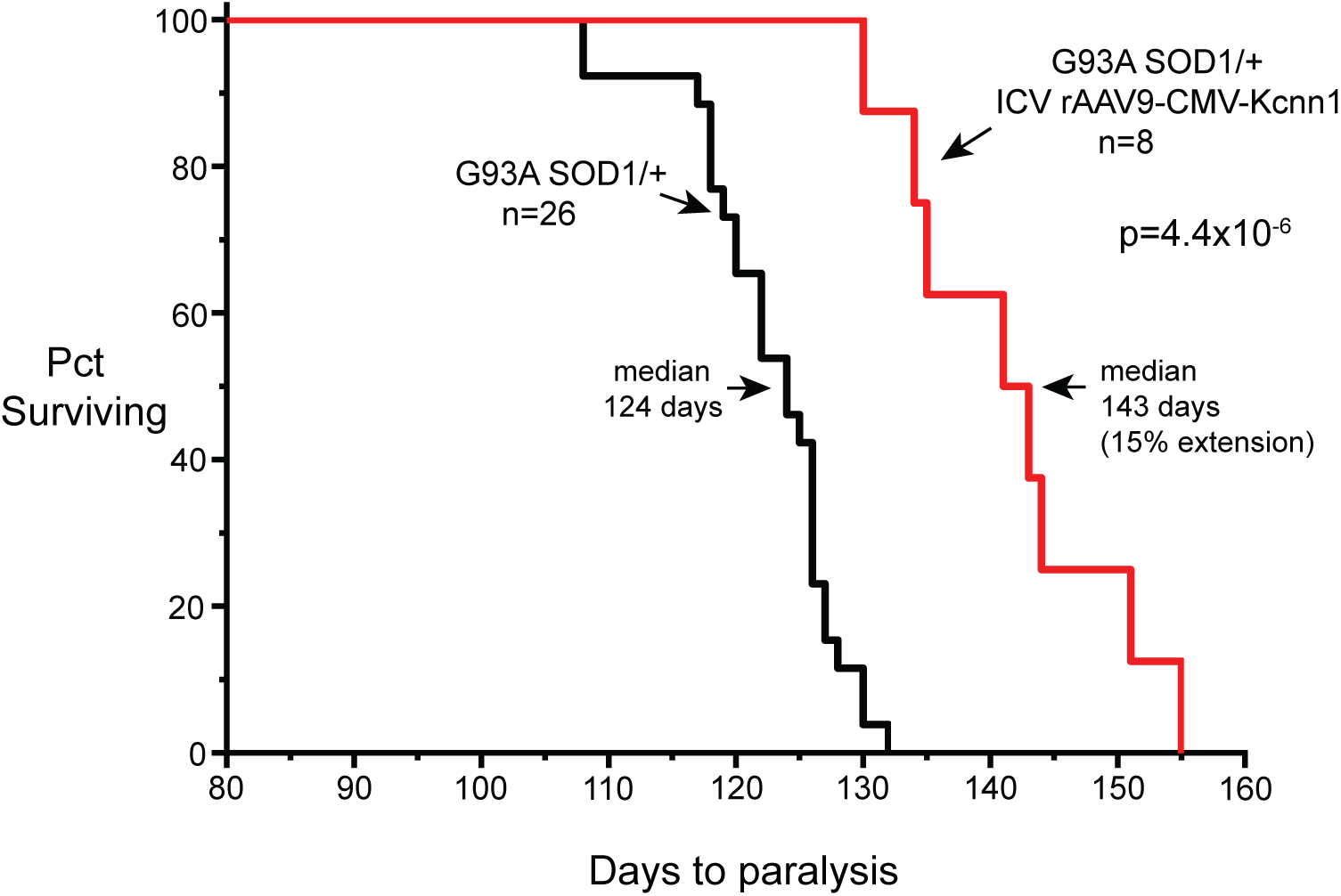
Kaplan-Meier survival plots of G93A/+ cohort and G93A/+ mice that had been injected at P0 into both lateral cerebral ventricles with self-complementary recombinant AAV9 carrying a CMV promoter driving mouse Kcnn1 cDNA, at titer of 5 × 10E13 vg/ml. Median survival to paralysis was increased by 15%. Notably, an earlier test using a titer of 1 × 10E13 produced only a few days of extended survival. See also SF8 and SF9 for RNA and immunostaining analysis of spinal cord from virus-injected mice.

### Thy1.2-Kcnn1 transgenic overexpression also extends the lifespan of A53T human alpha-synuclein transgenic mice

Could the effects of Kcnn1 transgenesis that extend survival and prevent aggregation in mutant SOD1 transgenic mice be of more general benefit? To address this question, we turned to a strain of A53T mutant human alpha-synuclein transgenic mice. Mutant alpha-synuclein shares with mutant SOD1 the property of misfolding and aggregation. Thy1.2 promoter-driven A53T synuclein transgenic mice (in a C57BL/6 background) have been reported to develop progressive lethal motor disease consisting of progressive rigidity/bradykinesia, and postural instability (Chandra et al., 2005; Martin et al, 2014) and, in our locally-carried strain, sometimes there has also been extremity paralysis. The clinical features have been associated with presence of aggregating human alpha-synuclein in cortex, hippocampus, basal ganglia, cerebellum, and brainstem (Martin et al, 2014). Because our locally-carried strain exhibited significant variability of clinical disease onset and only modest penetrance, we elected to move the A53T transgene from the C57BL/6 background into a B6SJL background (in which the various Thy1.2-Kcnn1 transgenes lie). The Thy1.2-A53T line was crossed to SJL/J mice, and indeed resulted in a strain with complete penetrance and lethality from motor symptoms with a median of 257 days (SF10). Notably, in this background, the motor symptom of rigidity was absent, bradykinesia was reduced, and later symptoms of ataxia and limb malpositioning/disordered gait became more prominent, with an endstage behavior of inability to maintain an upright position and consequently inability to locomote (SF11). We then mated the transgenic Thy1.2-A53T/B6SJL mice to Thy1.2- Kcnn1-3 to produce double heterozygous Kcnn1-3/+;A53T/+ mice. These exhibited a median survival time of 392 days, a 45% extension of survival (Fig.7). When the strain Kcnn1-6 was mated to the A53T/B6SJL strain, in an ongoing study, 4 of 6 mice have survived beyond 550 days, a time which is more than double the median survival time of A53T/B6SJL mice (Fig.7).

**Figure 7.**
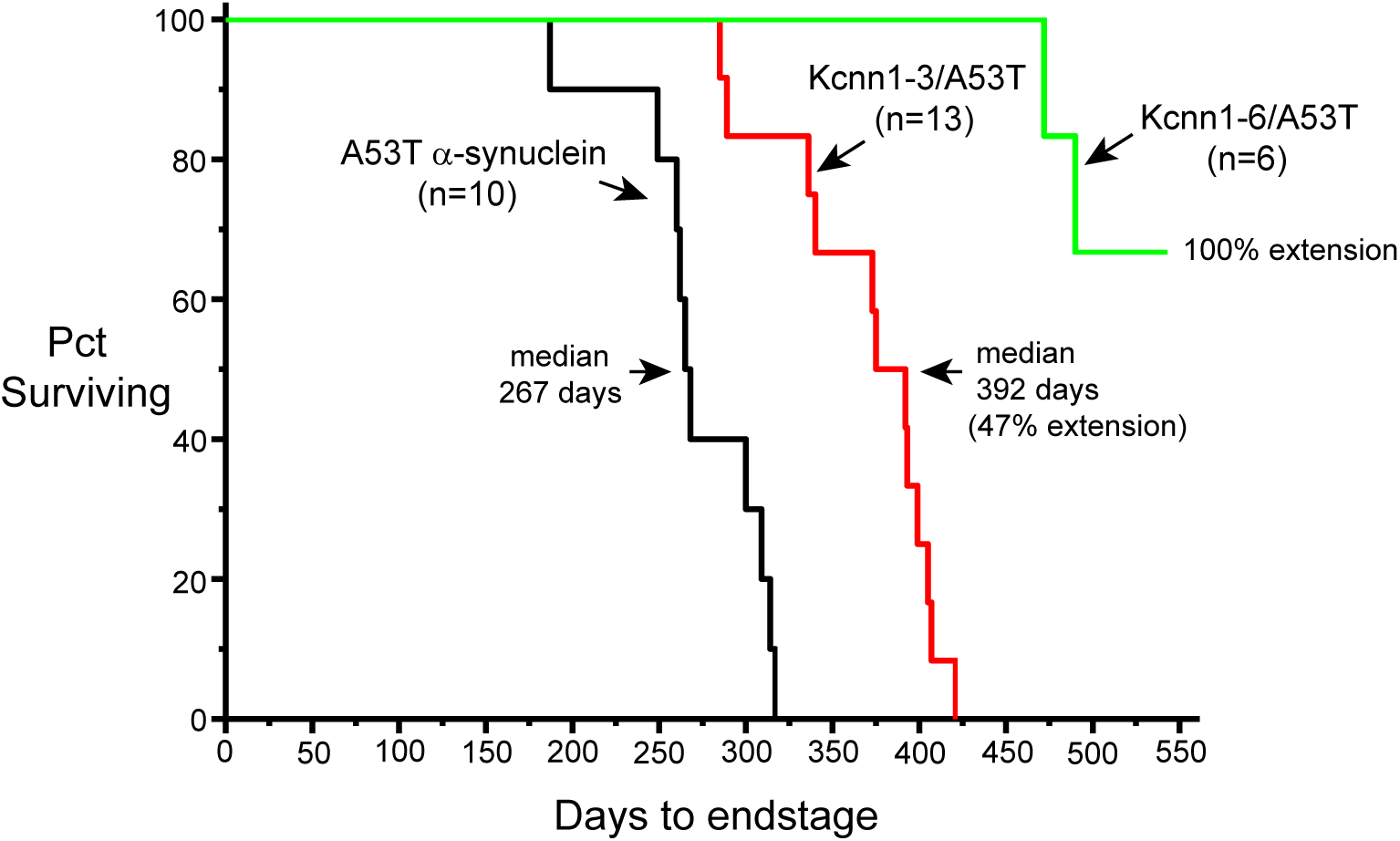
Kaplan-Meier survival plots of Thy1.2-A53T alpha-synuclein transgenic mice in a B6SJL background. Survival time is measured as days to the time of reaching an endstage motor behavior in which mice were no longer able to ambulate and could not maintain an upright stance (see SF11). The presence of Kcnn1-3/+ extended survival time by 47%, while presence of Kcnn1-6 (higher copy number) in an ongoing experiment has extended survival time by more than 100%.

To exclude an effect at the transcription level, e.g. involving suppression of A53T alpha-synuclein transcripts by Kcnn1 overexpression, qRTPCR of alpha-synuclein RNA was performed on brain RNA from A53T/+ mice, from A53T/+;Kcnn1-3/+ mice, and from A53T/+;Kcnn1-6/+ mice. There was substantial variability between individual mice of each strain, but no consistent relative depression of alpha-synuclein RNA in the presence of Kcnn1 (SF12).

As a marker of alpha-synuclein aggregation disease, we immunostained for the presence of S129-phosphorylated alpha synuclein (Karampetsou et al, 2017; Ghanem et al, 2022.). In A53T/B6SJL mice, we observed a small amount of this species in motor cortex at 7 months of age. In contrast, at endstage (9.5 months), we observed bright immunostaining in motor cortex, zona incerta, and deep cerebellar nuclei, and lesser staining in striatum, thalamus, subthalamic nucleus, and substantia nigra (Fig.8). We next examined a one year old Kcnn1-6/+;A53T/+ mouse, a time at which none of these mice have reached endstage. No S129-phosphorylated alpha synuclein could be observed, suggesting that the S129-phosphorylation pathologic process is not present. It thus appears that Kcnn1 overexpression exhibits a copy number-dependent beneficial effect on A53T alpha-synuclein-mediated pathology and survival, resembling the lack of aggregation and improved survival with increasing Kcnn1 transgenic copy level on mutant SOD1-linked disease.

**Figure 8.**
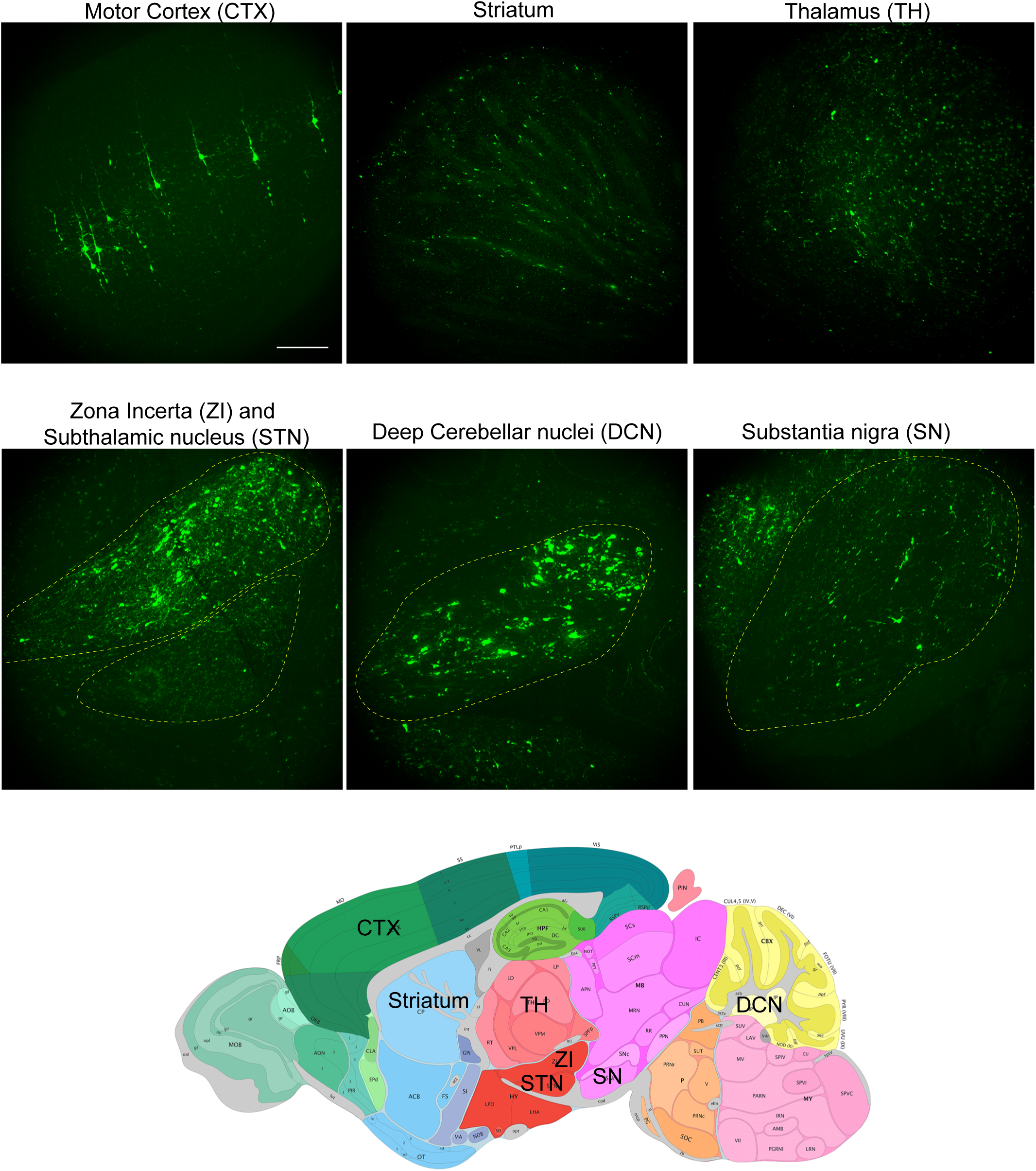
Figure 8. Anti-phosphorylated-S129 alpha-synuclein staining of sagittal sections of an endstage A53T alpha-synuclein/B6SJL mouse brain, showing intense signals in motor cortex, likely localizing to layer V, zona incerta, and deep cerebellar nuclei, with lesser strength signals in striatum, thalamus, and substantia nigra. A schematic of a sagittal brain section is shown below (from Allen Reference Atlas-Mouse Brain, atlas.brain-map.org).

### Assessing the beneficial effect of Kcnn1 overexpression: no major interference with transcription of mutant SOD1 or A53T alpha-synuclein

How could Kcnn1 overexpression be providing neuronal protection? One possibility is that overexpression reduces the transcription of the offending proteotoxic species. Transcriptional studies of both transgenic Kcnn1-6/G93A SOD1 mice (spinal cord ventral horns) and Kcnn1-6/A53T alpha-synuclein mice (brain) indicated that transcript levels of the respective G85R SOD1YFP and A53T alpha-synuclein species were not significantly reduced (Fig.9 and SF12).

**Figure 9.**
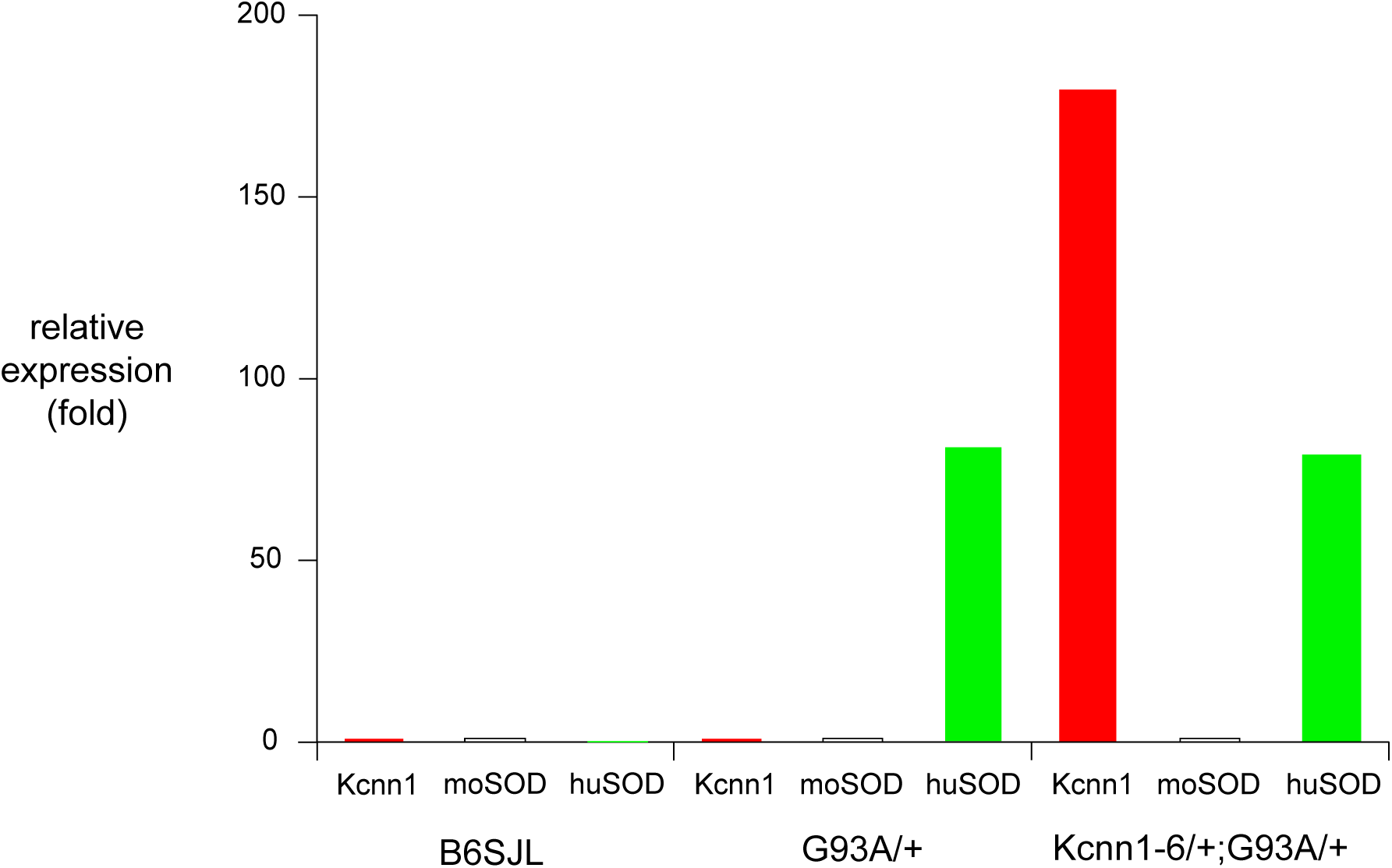
Kcnn1 overexpression in Kcnn1-6/+ spinal cord ventral horns does not affect the transcriptional expression of G93A human SOD1. Approximately 100 ventral horns of 20 micron thickness were laser capture-microdissected from each of B6SJL, G93A/+, and Kcnn1-6/+;G93A/+. RNA was prepared, and qRT-PCR performed on ∼10 ng of each RNA sample. The level of each indicated RNA was taken relative to an internal GAPDH reference and expression then measured as the fold difference with, in each sample, the B6SJL measurement as the control. Thus, for B6SJL, the values for Kcnn1 and mouse SOD1 were taken as 1. The presence of mouse Kcnn1-6 thus produced an ∼175-fold increase of Kcnn1 over endogenous mouse Kcnn1. For human SOD1 in G93A/+ and in Kcnn1-6;G93A/+, the ratio had to be taken relative to mouse SOD1 (since human SOD1 is absent from mouse). This ratio resulted in a value of ∼75 regardless of the presence of Kcnn1 transgene, indicating that expression of Kcnn1 does not impair the transcription of the pathogenic G93A transgene.

### A morphologic disturbance in spinal cord motor neurons of Kcnn1 transgenic mice observed by transmission EM suggests a stress response

The broad physical distribution of Kcnn1-encoded subunits in the cytosol of motor neurons of Kcnn1 transgenic mice observed by immunofluorescence suggested that it might be valuable to obtain higher magnification images of the motor neurons, e.g. to inspect for disturbance of organelle morphology. We thus prepared transmission EM samples of spinal cord motor neurons from perfused 3-4 month old Kcnn1-3/Kcnn1-3 mice, Kcnn1-6 mice, and B6/SJL mice. The EM images of motor neurons from both Kcnn1 transgenic strains were striking for the presence of an organellar disturbance as compared with B6SJL. Many cells exhibited expanded ER, in some cells manifesting as tight stacks of parallel ER cisternae, but most strikingly in some cells presenting as a whorl pattern of such parallel cisternae (Fig.10). In some cases, ribosomes studded the cisternae, but in others the ER appeared smooth. This seemed indicative of an ER stress response (Bommiasamy et al, 2009; Schuck et al, 2009).

**Figure 10.**
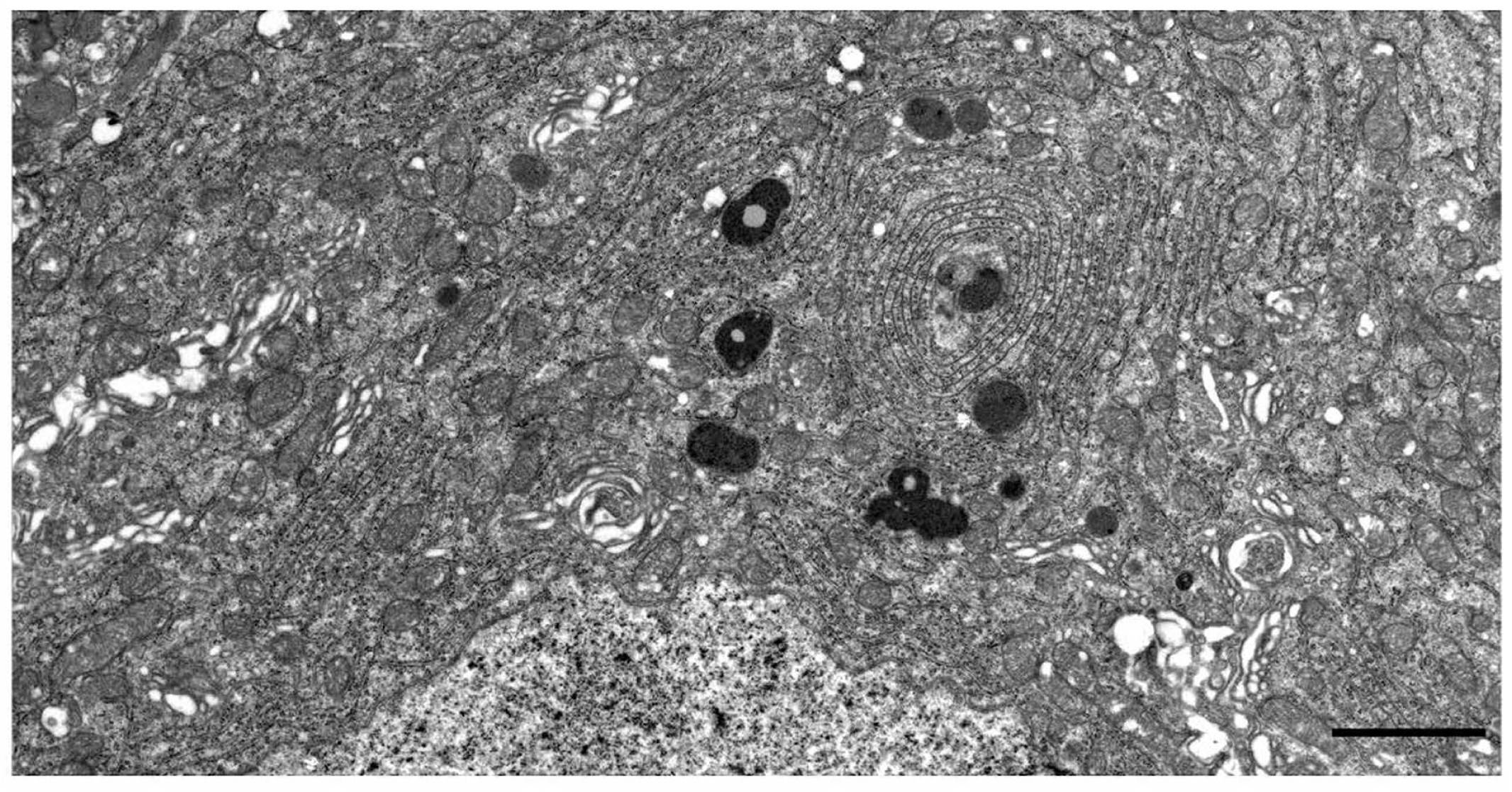
Expanded ER in spinal cord motor neuron of a 4 month old Kcnn1-6 mouse. Ultrathin sections were prepared following PFA perfusion, dissection of the spinal cord, PFA/glutaraldehyde post-fixation, epon embedding, and microtome sectioning. Note the whorl patterns of stacked ER membranes that tails off in a downward leftward direction. Darkly stained structures are lipofuscin granules, some containing typical lipid droplets. More lightly staining round structures are likely to be lysosomes. The nucleus is positioned at the bottom aspect of the image and surrounding Golgi has a dispersed appearance. Scale bar, 2 microns.

### RNAseq profiling of laser capture-microdissected motor neurons from Kcnn1-6 and B6SJL spinal cords reveals an ER stress response and a mitochondrial stress response in the Kcnn1-6 motor neurons

To assess for a Kcnn1-mediated stress response, RNAseq was carried out on RNA from pools of 1000 laser capture-microdissected spinal cord motor neuron somata from each of four B6SJL (wt) and four Kcnn1-6 transgenic mice. An average of 67 million paired reads were obtained from each sample. Reads were aligned to the mouse genome assembly (mm10), identifying ∼21,000 genes. DEG analysis was performed using DESeq2, and p-values for the differences were adjusted for multiple comparisons to give p-adjust (p_adj_) values. Significantly differentially expressed genes (p_adj_ <0.05; n=1788: 968 down, 820 up) were assessed for functional enrichments using Metascape. Of the 20 most significantly enriched categories in the gene ontology analysis, only two showed strongly upregulated transcription in Kcnn1-6 transgenics: GO:0034976 “Response to ER stress” and KEGG pathway mmu04141, “Protein processing in the ER” (see Fig.11). Table I lists the two sets of upregulated ER genes, including all of those exhibiting 30% or greater increases, along with significance values (as well as several relevant genes with a 20% increase). The genes could be assigned to functional categories of ER stress (as in Travers et al, 2000), reflecting the presence of effects at multiple levels of ER physiology.

**Figure 11.**
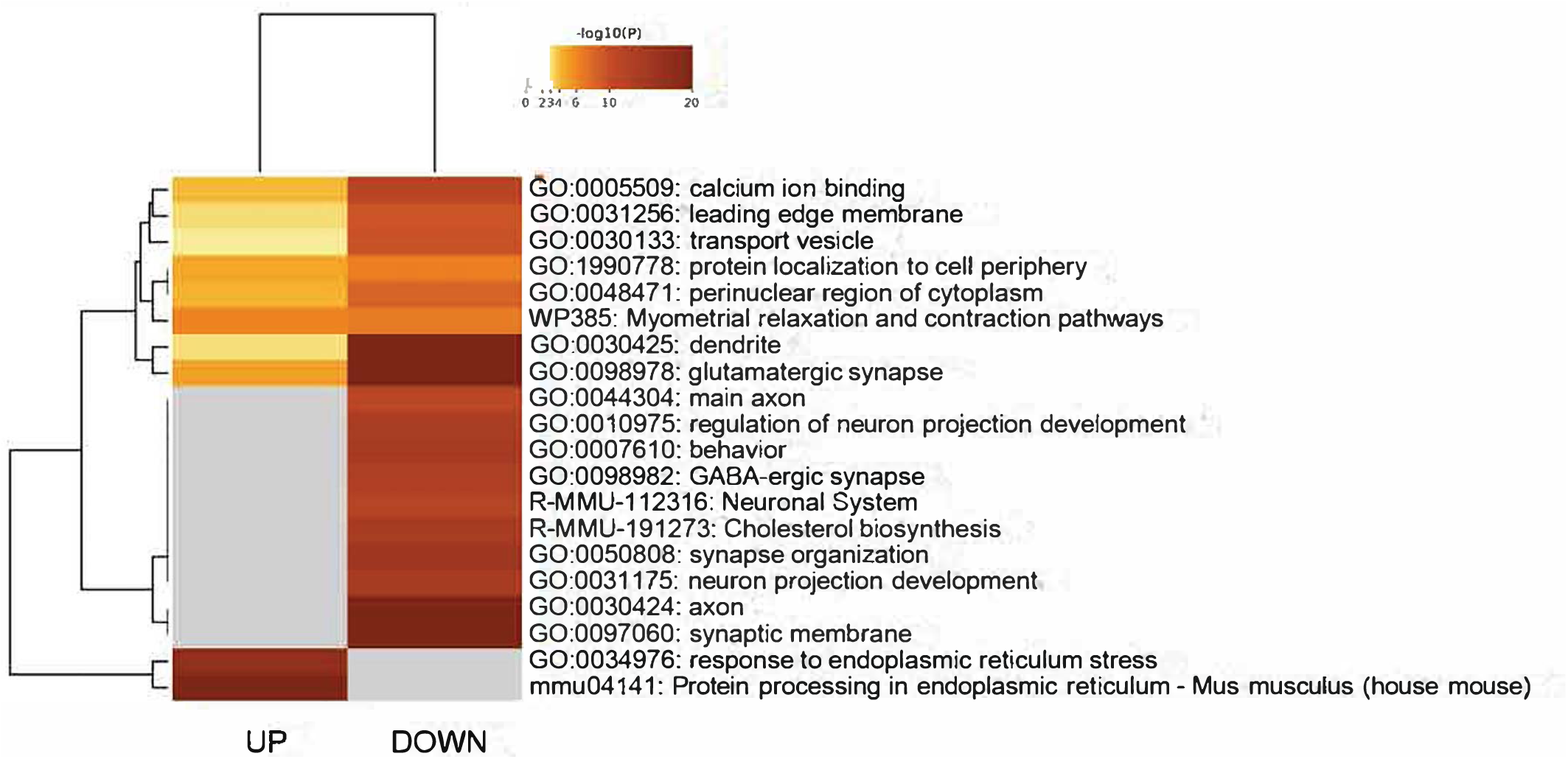
Heat map showing gene ontology analysis of RNAseq data comparing laser captured spinal cord motor neurons from B6SJL with those from Kcnn1-6 transgenic mice. Of the 20 most significantly enriched categories in the gene ontology analysis, only two showed strongly upregulated transcription in Kcnn1-6 transgenic mice, both involving the ER. (see text).

**Table I.** ER STRESS RESPONSE GENES UPREGULATED BY KCNN1 TRANSGENESIS.

|  | <u>Fold change</u> | <u>p adj</u> |
| --- | --- | --- |
| **STRESS SIGNALING** |  |  |
| Creb3L1 | 4.0 X | 2 x 10E-5 |
| GADD34 (Ppp1r15a) | 1.4 X | 4.5 x 10E-3 |
| XBP1 | 1.3 X | 6.2 x 10E-3 |
| ATF4 | 1.3 X | 2.4 x 10E-2 |
| CHOP (Ddit3) | 1.6 X | 2.9 x 10E-7 |
| Trib3 | 3.0 X | 1.4 x 10E-2 |
| Sestrin2 | 1.3 X | 5.4 x 10E-5 |
| ASNS | 1.2 X | 2.5 x 10E-2 |
| ATF5 | 2.7 X | 4.5 x 10E-13 |
| ATF6 | 1.2 X | 1.2 x 10E-3 |
| WFS1 | 1.5 X | 1.3 x 10E-6 |
| **TRANSLOCATION** |  |  |
| Sec61 beta | 1.6 X | 3 x 10E-8 |
| Sec61 gamma | 1.6 X | 2.2 x 10E-9 |
| TRAM | 1.5 X | 1.2 x 10E-7 |
| Ssr1 (signal seq R) | 1.3 X | 4 x 10E-7 |
| Ssr4 (signal seq R) | 1.3 X | 5 x 10E-4 |
| SERP1 | 1.4 X | 2.4 x 10E-8 |
| **PROTEIN FOLDING** |  |  |
| **Chaperones** |  |  |
| BiP (HspA5) | 1.5 X | 9.7 x 10E-13 |
| DnaJB9 | 1.5 X | 6.8 x 10E-12 |
| GRP94 | 1.3 X | 5.2 x 10E-5 |
| CryAB | 1.6 X | 9.2 x 10E-11 |
| HYOU (ER Hsp110) | 1.5 X | 3 x 10E-15 |
| Calnexin | 1.2 X | 2.2 x 10E-4 |
| Glucosidase 2 (Prkcsh) | 1.3 X | 5 x 10E-4 |
| **Protein disulfide isomerases** |  |  |
| PdiA3 | 1.2 X | 1.6 x 10E-3 |
| PdiA4 | 1.3 X | 1 x 10E-2 |
| PdiA6 | 1.6 X | 4 x 10E-9 |
| P4hb (disulfide isom/prolyl4OHase) | 1.5 X | 1.6 x 10E-17 |
| **GLYCOSYLATION/MODIFICATION** |  |  |
| Ostc (oligosacch.xferase subunit) | 1.6 X | 1.6 x 10E-10 |
| Magt1 (oligosacch xferase subunit) | 1.9 X | 3.8 x 10E-10 |
| TMEM258 (oligosacch xf subunit) | 1.4 X | 3.3 x 10E-4 |
| Lman (ERGIC 53) | 1.3 X | 3 x 10E-3 |
| GRASP55 | 1.3 X | 2.7 x 10E-5 |
| **ER-ASSOCIATED DEGRADATION** |  |  |
| Derl3 | 3.4 X | 1.0 x 10E-2 |
| Sdf2l1 | 2.0 X | 6.0 x 10E-12 |
| Herpud1 | 1.3 X | 3.9 x 10E-5 |
| Edem2 | 1.3 X | 3 x 10E-4 |
| FBXO6 (FBS2) | 1.4 X | 2.4 x 10E-2 |

Under “Stress Signaling”, XBP1 mRNA was modestly induced, but more revealing, the activating splice of XBP1 RNA was observed in 36% of the Kcnn1-6 reads and in none of the B6SJL reads. This result implies that the IRE1 limb of the UPR is activated in Kcnn1-6 motor neurons (Walter and Ron, 2011), contributing e.g. to induction of RNAs encoding ER chaperones (BiP/Hspa5 and cooperating component DnaJB9 as examples; see Shen et al, 2002). RNAs for components of the translocon were also induced, as well as for (ER-localized) protein disulfide isomerases and glycosylation components. Significantly, RNAs encoding ERAD components such as Derlin 3, a component of the HRD1 retrotranslocation complex known to be induced under ER stress (Eura et al 2020), were also induced. We conclude that there is a chronic and, lacking an analysis of distribution of the changes amongst 1000 motor neurons that were studied, at least modest degree of ER stress response in the spinal cord motor neurons of the Kcnn1-6 expressing mice, and that this may contribute to neuroprotection.

The likely trigger of the ER stress response is the Kcnn1 protein itself, which, like other potassium channels, is normally targeted to the ER for biogenesis (Papazian, 1999; see also Hegde and Keenan, 2024). Here, in an overproduced state, the Kcnn1 subunit likely at least mildly stresses the ER, presumably as the result of misfolding or misassembling of some fraction of the overexpressed Kcnn1-encoded subunits.

### An integrated stress response is also present

A low level of induction of ATF4 RNA was observed, as well as induction of RNAs of several of ATF4’s downstream transcriptional targets, CHOP (Ddit3), Trib3, Sestrin2 (Sesn2), and Asns (see Table I), raising a question of whether there might be activation of the integrated stress response (ISR). Hallmark features of an ISR include phosphorylation of Ser51 of eIF2alpha to activate the ISR (by any one of four different kinase sensors, including PERK (*EIF2AK3*), which recognizes unfolded/misfolded ER proteins). Such phosphorylation of eIF2alpha reduces translation initiation at a global scale, and is accompanied by translation of a select group of uORF-containing mRNAs, including that for ATF4 (see Costa-Mattioli and Walter, 2020). Indeed when we compared immunostained spinal cord cross-sections of B6SJL with Kcnn1-6, we observed increased levels of both phosphorylated eIF2alpha and ATF4 in the ventral horn (Fig. 12). In the former case, there was a several-fold increase of cytosolic signal in Kcnn1-6 motor neurons (Fig. 12A, left panel). In the case of ATF4 (Fig. 12B), in B6SJL motor neurons, there was a weak cytosolic signal (top row, right), while by contrast, in Kcnn1-6 motor neurons, the ATF4 signal was nuclear-localized and strong in some cells, but weaker in others (top row, left). We conclude that an ISR is present.

**Figure 12.**
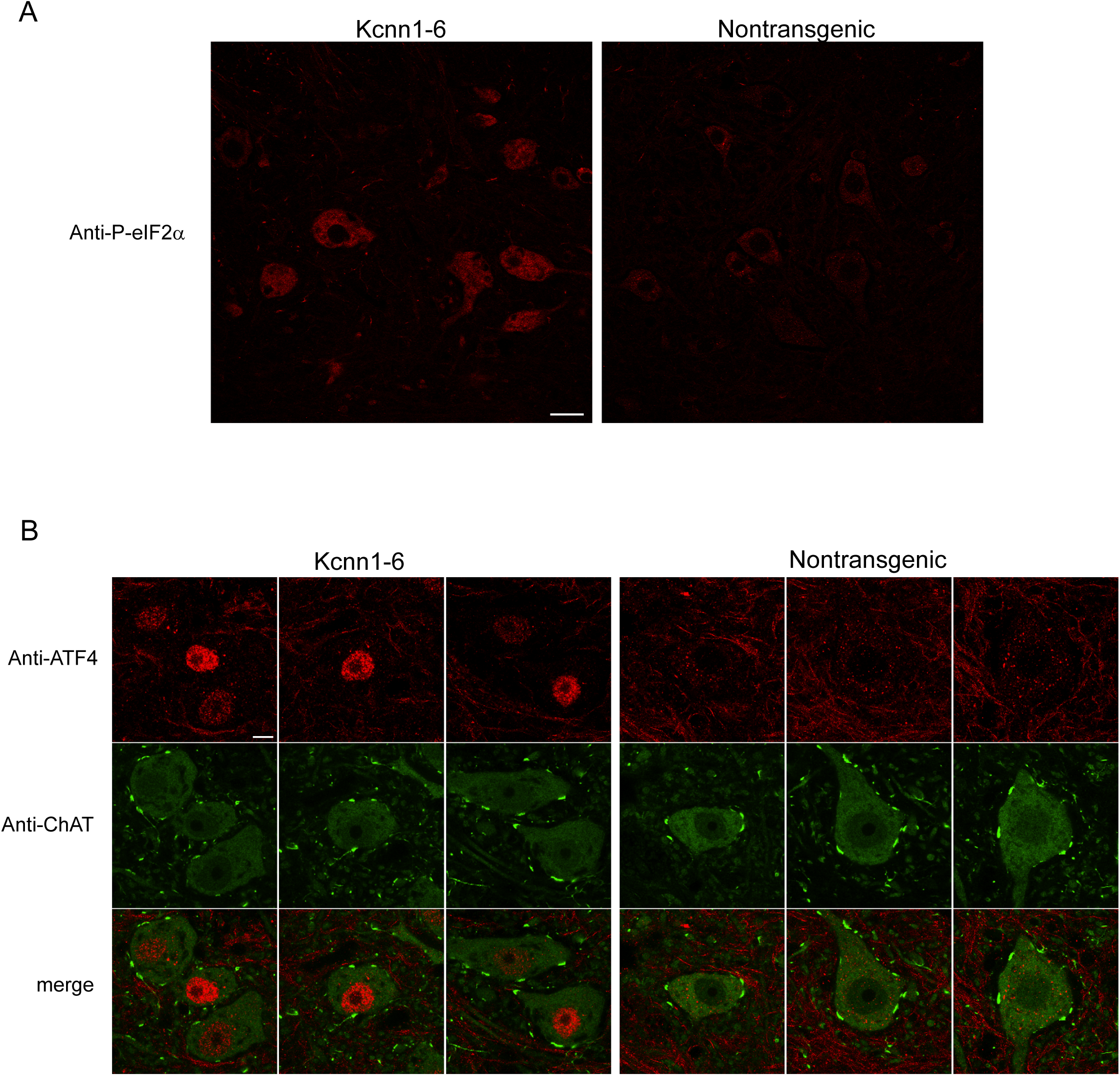
Presence of an integrated stress response (ISR) in Kcnn1-6 transgenic mice. Immunostaining with anti-phosphorylated-S51-eIF2alpha (A) and anti-ATF4 (B), comparing spinal cord sections of a Kcnn1-6 mouse with those of a nontransgenic B6SJL mouse. (A) shows an at least several-fold increase of anti-phospho-S51-eIF2alpha signal in Kcnn1-6 motor neurons (left) compared with that in those of the B6SJL mouse (right), reflecting activation of the ISR. (B) shows three representative ventral horn sections from a Kcnn1-6 mouse and three from a B6SJL mouse. Motor neurons are identified by anti-ChAT antibody staining (middle row of images). The Kcnn1-6 sections (upper left images, first and third with 3 and 2 motor neurons, respectively) show a variably intense anti-ATF4 signal that is localized to the nucleus. This compares to a relatively weak or absent signal that is localized to the cytoplasm in the nontransgenic cord (upper right images). This reflects the selective translation that occurs as a feature of the ISR. Scale bar 20 μm in (A), 5 μm in (B).

The effect of reduced translational initiation could extend to mutant SOD1 and A53T alpha-synuclein, lowering their overall levels, preventing aggregation/toxicity.

### Mitochondrial stress response

We also observed strong induction of RNAs implicated in a mitochondrial stress response (Table II)(see Melber and Haynes, 2018; Anderson and Haynes, 2020). This included the regulatory transcription factor ATF5 (which can be induced by ATF4 from the ISR pathway), as well as downstream factors induced by it, such as FGF21 and GDF15 (the latter two factors noted to be induced, for example, in children with primary mitochondrial diseases, Riley et al, 2021). The partitioning between nucleus and mitochondria of the regulatory factor, ATF5, may be affected here to favor ATF5 nuclear localization and induction of a mitochondrial stress response (also termed UPR^mt^).

**Table II.** MITOCHONDRIAL STRESS RESPONSE GENES UPREGULATED BY KCNN1.

|  | **<u>Fold change</u>** | **<u>P adj</u>** |
| --- | --- | --- |
| FGF21 | 36 X | 1.2 x 10E-12 |
| GDF15 | 11.6 X | 8.3 x 10E-5 |
| Atf5 | 2.7 X | 4.5 x 10E-13 |
| Aifm2 | 1.8 X | 6 x 10E-8 |
| MAVS | 1.7 X | 2 x 10E-2 |

### Additional transcriptional responses

In a further possible chaperone response in the cytosol, RNA encoding the Hsp70 protein, Hsp1a1, was also upregulated in the Kcnn1 transgenic motor neurons (1.56 X; p=0.035). RNA encoding the integral membrane autophagy component, ATG9b (Chiduza et al., 2023), was also induced (1.4X, padj = 1 × 10E-4). A preliminary immunostaining with anti-phospho S757 ULK1 antibodies of spinal cord sections observed reduced signal in Kcnn1-6 relative to B6SJL, also supporting upregulation of autophagy. There was also evidence for an oxidative stress response, with RNA for aldehyde dehydrogenases Aldh1a2 and Aldh1a7 induced 4.3-fold and 5-fold, respectively (padj = 0.01 and 0.03; see Singh et al, 2013).

## Discussion

We have reported here that Thy1.2 promoter-driven neuronal overexpression of the mouse calcium-activated small conductance potassium channel subunit Kcnn1 is associated with extended survival of transgenic mutant G85R and G93A SOD1 ALS mouse models and a transgenic A53T mutant alpha-synuclein mouse model. The disease models feature overproduction of two distinct cytosolic proteins (mutant SOD1 and mutant alpha-synuclein), characteristically misfolded and aggregated in affected neuronal tissues of patients. Here, the neuronal Thy1.2 promoter-driven expression of Kcnn1 cDNA at the level of 3-8 transgene copies (by qPCR of tail DNA but probably greater number based on genomic sequencing) was associated with 5-10 fold overexpression of the Kcnn1-encoded subunit observed by immunostaining of motor neurons and was able to substantially prolong survival of both mouse models (Figs1,2,6 and SF3). How is overexpression of Kcnn1 able to mediate such effects?

### Action of overexpressed Kcnn1

There are two major mechanisms that could explain the observed favorable effects on survival, one involving electrophysiology and the other involving induction of a neuronal stress response. By the former mechanism, overexpression of Kcnn1-encoded subunits could result in an increased number of putative Kcnn1 subunit-containing tetrameric channels. This could prolong hyperpolarization and decrease overall neuronal firing rate, potentially relieving energetic/metabolic stress on neurons already coping with a misfolded neurotoxic protein.

Yet in the case of the rodent Kcnn1-encoded subunit, used in the present study, there is evidence indicating its inability to form active SK1 channels in the setting of overexpression. A study reporting on rat Kcnn1-encoded subunit overexpressed in HEK293 cells indicated that the subunits alone could not form an active channel that would conduct a calcium-dependent potassium current (Benton et al, 2003, Fig.1). By contrast, rat Kcnn2, which homo-oligomerizes, readily produced calcium-dependent potassium current. Only when rat Kcnn1 was co-expressed with rat Kcnn2 was any effect of Kcnn1 observed, with Kcnn1 increasing the current measured for any given constant voltage condition. This was taken as evidence that rat Kcnn1 achieves activity only by hetero-oligomerizing with rat Kcnn2. Whether the Kcnn1-encoded subunit behaves, when expressed alone, as a monomer or as an inactive tetramer has not been directly resolved. However, Benton et al observed a patchy cytoplasmic pattern of the overexpressed protein and absence of cell surface localization in their Kcnn1-transfected cultured HEK cells (Figure 2 therein). This pattern is similar to that which we observed here in motor neurons of mouse Kcnn1-overexpressing transgenic mice (Fig.4). The intracellular distribution of Kcnn1 in both systems is consistent with failure of biogenesis to be completed, suggesting that overexpressed rodent Kcnn1 likely remains as an unassembled and possibly misfolded monomer. The monomeric states suggested by this model would likely induce stress responses in motor neurons as reported here in Figs.10-12 and Tables I and II.

Consistent with the foregoing proposed model, Thy1.2-driven overexpression of mouse Kcnn2 did not produce any beneficial effect in our survival assay of homozygous G85R SOD1YFP (SF4), even at copy numbers 10-fold greater than any of the Thy1.2-Kcnn1 transgenics. We interpret such an outcome as indicating that overexpressed Kcnn2 homotetramerized and formed an active channel, as in the overexpressing HEK cells, and, while that may have affected AHP, it did not confer protection from mutant SOD1.

Overall, we surmise that overexpressed mouse Kcnn1 is targeted, like other newly-synthesized potassium channel subunits, to the ER (Papazian, 1999), but in the absence of high levels of Kcnn2 with which to coassemble, the subunit persists as either misfolded or misassembled Kcnn1 species. These species produce, as primary responses directed from the ER, an ER stress response and an integrated stress response (the latter presumably triggered by misfolding of Kcnn1, potentially activating the transmembrane PERK kinase). PERK could, via phosphorylated eIF2alpha inhibition of general translation, reduce translation of the pathogenic proteins, mutant SOD1 and A53T synuclein, reducing toxicity/aggregation and improving survival.

The basis to a mitochondrial stress response and induction of other stress-responsive pathways, potentially including autophagy (which could also offer a removal pathway for the pathogenic SOD1 and synuclein species), is less clear. Perhaps simple induction of ATF5 via ATF4 can lead to the mitochondrial stress response. In sum, it seems like mouse Kcnn1 overexpression is able to prime against proteotoxicity by mutant SOD1 and A53T synuclein neuropathogenic species.

It remains to be seen whether the chronic stress program observed here can be instituted at later times than the one or two weeks postnatal when the Thy1.2 promoter becomes active, e.g. at times when motor symptoms are developing in the affected mice. It also remains unknown whether the program observed here could counter other cytoplasmic pathogenic species such as tau or TDP43.

### Clinical effects on mice of Thy1.2-Kcnn1 expression alone

The beneficial effects of transgenic Kcnn1-3/+ were without any visible additional effects. That is, Kcnn1-3 mice were normal in size, mobility, and behavior, including breeding. In fact, the Kcnn1-3 mice may be especially long-lived. Three mice that were set aside during early experiments have survived to near or beyond 3 years of age. In particular, one mouse is now 1100 days old (3 yr = 1095 days); one mouse died of abdominal tumor at 958 days, 5 months short of 3 years; and one mouse is now 950 days old and in good health, 5 months short of 3 years. Additional Kcnn1-3/+ mice as well as control B6SJL mice are being monitored (we note that several of the control mice have died by 15 months of age). This said, the higher copy Thy1.2-Kcnn1-6 transgenic mice developed mild lower extremity spasticity at 4-6 months of age (not present in any of the Kcnn1-3/+ mice), but it did not progress, and these mice are in many cases now 2 years old.

### Further studies/implications

Clearly it will be important to resolve the precise localization(s) of Kcnn1 in the motor neurons of the transgenic strains. In further localization tests, Kcnn1 has seemed to follow a cellular distribution that corresponds to the ER but super-resolution studies will be needed to resolve whether there is precise overlap. One of the difficulties is that the anti-Kcnn1 antibody is only functional in postfixation conditions, preventing normal PFA perfusion preparation of tissue. However, it seems that better resolution of the location of the protein, clarification of whether it is a monomer or a superstructure, and whether it has bound calmodulin (which could serve as a conformational stabilizer or calcium sink) should all help to better define how a multi-faceted stress response arises and thus define the nature of the protective action. We also do not know how pathogenic cytosolic protein is being removed vs e.g. simply being bound, as a protective mechanism. “Removal” could effectively involve any of reduced translation, autophagy, proteasomal degradation, or exocytosis. Finally, there may be implications of the AAV-mediated Kcnn1 transduction experiments (Fig.5 and SF 8,9), suggesting that rAAV delivery to susceptible neurons could confer clinical benefit.

## Acknowledgments

This work was supported by HHMI, a Yale Sterling Endowment, and AWM Bio Inc. We thank the Yale Animal Resources Center and the Yale Genome Editing Center for superb technical assistance. We thank Sreeganga Chandra for helpful discussions. A.L.H., W.A.F., and M.N. are co-founders of AWM Bio, Inc. which is the exclusive licensee of certain pending patent applications covering this work.

## Methods

### Mice

All animal experiments were carried out under protocols approved by the Yale University Institutional Animal Care and Use Committee in accordance with National Institutes of Health guidelines for the ethical treatment of animals. The transgenic G85R SOD1YFP mouse strain, which has a human genomic SOD1 sequence including the promoter, the codon 85 GGC to CGC change in exon 4, and the addition of the YFP coding sequence to the last SOD1 codon in exon 5, has been described previously (Wang et al., 2009; Hadzipasic et al., 2014). Mice heterozygous for a transgene of Thy1.2-human A53T alpha-synuclein cDNA were from Chandra et al. (2005) and had been maintained in a C57BL6 background. Transgenic G93A mutant SOD1 mice (JAX strain #002726), Chat-EGFP mice (JAX strain #007902), and wild-type B6SJLF1/J mice, here called B6SJL (JAX strain #100012) were purchased from Jackson Laboratories. Transgenic and CRISPR/Cas knockout lines were produced by the Yale Genome Editing Center.

All animals were genotyped by quantitative real-time PCR using primers suggested either by PrimerBank (pga.mgh.harvard.edu/primerbank) or by the primer designing tool Primer-BLAST (www.ncbi.nlm.nih.gov/tools/primer-blast/) from NCBI. All primer sets were tested to ensure linearity and efficiency >90%. Specificity was confirmed by sequencing the amplified products.

### Transgenic Constructs

All constructs used to produce the transgenic strains were based on the Thy1.2 vector, a gift from P. Caroni, Friedrich Miescher Institute, Basel (Caroni, 1997), modified by insertion of a short double-stranded oligonucleotide into the XhoI site to permit directional cloning between XhoI (5’) and AgeI or SalI (3’) sites. Plasmids containing the following mouse cDNA clones were obtained from OriGene Technologies: Kcnd2, MR206092; Gabra1, MR227303; Chrm1, MR207339; Chrna4, MR209587; and Kcnn1, MR208601. Mouse Kcnj2 was from Addgene plasmid 60598. For all constructs, PCR amplification was used to recover the cDNA sequences without any tags and with appropriate terminal restriction sites to allow cloning into the Thy1.2 vector. All constructs were sequenced after preparation. Restriction enzyme digestion was used to release the complete transgenesis vector, which was subsequently purified by agarose gel electrophoresis.

### Viruses

A plasmid [pscAAV-GFP, Addgene plasmid #32396, a gift from John T. Gray, St. Jude Children’s Research Hospital, Memphis, TN (Gray et al., 2011)] to produce self-complementary AAV-CMVGFP virus was modified by direct replacement of the GFP sequence with one encoding ECFP (Clontech) to produce pscAAV-CFP. The plasmid psc-CMV-Kcnn1 was similarly produced from the same starting plasmid by inserting the mouse Kcnn1 cDNA sequence (NM_032397), PCR-amplified from OriGene Technologies plasmid MR208601, in place of GFP, omitting the 3’-terminal Myc-DDK tag sequence. These plasmids were packaged at the University of North Carolina Chapel Hill Vector Core to produce recombinant AAV viruses with the AAV9 serotype at titers between 1.6 x 10^13^ and 4.5 x 10^13^ vp/ml.

### P0 ICV Injections of AAVs

Female B6SJL mice were bred with male G93A SOD1 mice, and their pregnancies were closely followed. Shortly after birth (4-8 hours), pups were examined for visible milk in their stomachs and, if present, then used for injection. The general injection protocol closely follows Kim, J.-Y., et al. JoVE 91: e51863, 2014. A digitally controlled, motorized stereotactic apparatus (Stoelting) was used for anesthetizing the animal by hypothermia, restraining it with padded ear bars, positioning the needle, and carrying out the injection. A 10 µl Hamilton syringe loaded with virus was placed in the vertical holder on the apparatus. The needle was positioned at lambda, and the (x,y) stereotactic coordinates of lambda in mm were set as (0,0) on the digital micrometer. Then the position of the needle was changed to (0.8,1.5), which placed it halfway from lambda to bregma and 0.8 mm lateral to the sagittal suture. The needle was lowered until it just penetrated the skin, set as z = 0, and then lowered an additional 1.7 mm to achieve penetration of the skull and ventricle. After backing the needle out to 1.5 mm, 1.5 µl of virus solution was slowly injected mechanically over ∼90 sec. The needle was left in place for 60 sec, then slowly backed out completely (1-2 min). The apparatus was adjusted to (-0.8,1.5), which placed it over the contralateral ventricle, and the injection was repeated on the other side. After injection, anesthesia was reversed by placing the pup on a warming pad, and the pups were returned to their dam and allowed to mature normally. Pups were genotyped before weaning to distinguish the mutant G93A animals from their non-transgenic littermates, and both were retained for observation of time of survival.

### qRT-PCR Transcript Analysis

Transcripts for SOD1, Kcnn1, and α-synuclein were quantitated by qRT-PCR following RNA preparation from one hemisphere of a brain, an entire spinal cord, a 3 mm section of spinal cord, or a collection of spinal cord ventral horns, in all cases taken under RNase-free conditions from a transcardially PBS-perfused mouse and flash-frozen in LN_2_. Brain and whole spinal cord were crushed in a LN_2_-cooled stainless steel mortar and pestle; the powder was suspended in RLT+ buffer (Qiagen) then homogenized with a TissueRuptor (Qiagen). The frozen sections of cord were suspended in RLT+ and homogenized directly. RNA was purified from about 25% of the brain homogenate and all of the spinal cord homogenate using a RNeasy Plus Mini Kit (Qiagen); RNA from the cord sections was purified with a RNeasy Plus Micro Kit (Qiagen). For analysis of RNA in motor neuron-rich parts of the spinal cord, ∼100 ventral horns were laser microdissected from spinal cord sections directly into RLT+ buffer, and the RNA was purified with a RNeasy Plus Micro Kit. Transcripts were converted to first strand cDNAs with the PrimeScript 1st strand cDNA Synthesis Kit (Takara) using a mixture of the supplied oligo(dT) and random primers (1:2, respectively). The cDNAs were quantitated directly by real-time PCR using previously validated primers. Mouse GAPDH and mouse SOD1 primers were used as internal reference controls and for ΔΔC_T_ calculations.

### Antibodies

Antibodies used: anti-mouse Kcnn1 (Proteintech 17929-1-AP, 1:400), anti-ATF4 (Cell Signaling 11815, 1:100), anti-CHAT (Abcam ab178850, 1:100), anti-phospho(S757) ULK1 (Cell Signaling 6888, 1:100), anti-phospho(S51) eIF2α (Cell Signaling 3398, 1:100), anti-phospho(S129) α-synuclein (Abcam ab51253,1:800), anti-Ctip2 (Abcam ab18465, 1:100).

### Immunostaining

Two preparations of mouse tissues were used, depending on whether immunostaining for Kcnn1 was being performed. When Kcnn1 immunostaining was not to be performed, mice were transcardially perfused with 4% (wt/vol) paraformaldehyde (PFA) in PBS, spinal cords and brains were dissected, and tissues were post-fixed overnight in 4% PFA. Following cryoprotection in 30% sucrose (>1 day), tissues were embedded in OCT (Tissue-Tek), frozen on dry ice, and stored at - 80°C. Alternatively, because of the requirements of the Kcnn1 antibody, when Kcnn1 staining was to be performed, mice were transcardially perfused with PBS alone. After dissection, brain segments and ∼3 mm cross-sections of spinal cords were immediately embedded in OCT and frozen on LN_2_-chilled methyl butane, then stored at −80°C. In both cases, 20 µm cross-sections of spinal cord and coronal or sagittal sections of brain were prepared on a cryostat (Leica CM3050S) and placed on Surgipath X-tra slides. Slides for Kcnn1 staining were post-fixed for 20 minutes in 4% paraformaldehyde in PBS, then washed 3 times in PBS before use. Blocking and staining were carried out as in Thomas et al., 2017.

### Image Acquisition

Most images were acquired on either a Leica SP5 or a Leica SP8 confocal microscope, using 40x or 63x oil objectives, as indicated. Laser lines appropriate for CFP, YFP, AlexaFluor 568, and AlexaFluor 647 were used for excitation, depending on the microscope and the fluor examined. Leica LAS software was used to display and export the images as TIF files for producing illustrations. Low power (10X) images were acquired on an Olympus IX81 microscope.

### RNA FISH

Stellaris RNA FISH was carried out on frozen sections of spinal cord or brain from B6SJL and Kcnn1-3 (homozygous) mice, prepared under RNase-free conditions. Probes were designed against the coding sequence of mouse Kcnn1 (NM_032397) using the Stellaris FISH Probe Designer (www.biosearchtech.com/stellarisdesigner) and labeled with CAL Fluor Red 610. Hybridization, washing, etc. were carried out essentially as described in the protocol for Stellaris RNA FISH provided by Biosearch Technologies (www.biosearchtech.com/stellarisprotocols). Imaging was performed as outlined above.

### Electron Microscopy

EM was carried out on 4% paraformaldehyde perfused and paraformaldehyde/glutaraldehyde post-fixed spinal cord segments of Kcnn1-6 mice. Preparation of ultra-thin (50 nm) sections and imaging were performed by the Yale Center for Cellular and Molecular Imaging (CCMI) Electron Microscopy Facility.

### RNAseq Analysis

RNA for RNAseq analysis was prepared from laser microdissected neuron cell bodies essentially as described in Bandyopadhyay et al. (2013). In all cases, transverse sections were prepared from OCT-embedded, flash-frozen tissues dissected from transcardially PBS-perfused animals under RNase-free conditions.

### Cranial nerves

Transverse sections of 20 micron thickness from brainstem/spinal cord of 2 month old ChAT-EGFP transgenic mice were cryosectioned one at a time from rostral-to-caudal, starting in the midbrain region above CN3, transferring to glass slides, and inspecting one at a time for GFP fluorescence until the CN3 nucleus was first observable. Thereafter, sections were transferred to PEN-membrane slides (Leica; one section per PEN slide) and the slides were kept at −80°C until preparation and laser capture. Briefly, preparation involved, one section (one slide) at a time, Azure B staining of the thawed slide, initial wash/fix in 70% ethanol to remove OCT, staining in 1% Azure B/70% ethanol (which identifies large motor neurons by staining the nucleus and cell body), followed by destaining in 70% ethanol (see Bandyopadhyay et al., 2013 for details). Laser capture was immediately carried out on the dried Azure B-stained slide with a Leica LMD6000 microscope, 20X objective. 20-30 dissected motor neurons were collected into 30 µl of RULT buffer in the cap of a 0.6 ml microfuge tube, which was briefly centrifuged then frozen at −80°C. A similar approach for CN12 was taken with the same tissue samples, carrying out further rostral-caudal dissection onto glass slides until CN12 in the medulla was first identified, then switching to transfer to PEN slides. Motor neurons were identified, laser dissected, and collected as for CN3 (above) from ∼50 sections. The collections of CN3 or CN12 neurons were then combined and RNA recovered by use of a Qiagen RNeasy UCP Micro Kit. Spinal cord motor neurons were similarly prepared, except that typically ∼10 cross-sections were applied to a single PEN slide.

RIN numbers were ∼7 for all samples. Libraries were prepared from 3-9 ng of each RNA using the NEBNext poly(A) mRNA Magnetic Isolation Module and the Ultra II RNA Library Prep Kit for Illumina with multiplex index primers. Libraries were subjected to quality control analysis on a Bioanalyzer DNA chip and were quantitated using the NEBNext Library Quant Kit. Libraries were diluted to 4 nM, pooled and denatured according to the instructions for Illumina NextSeq 550. Libraries were sequenced with settings for single-index, paired-end sequencing with 75 cycles per end. Three independent replicates each of of 12N, 3N, and spinal cord were sequenced. An average of 105 million paired-reads were obtained from each sample (min 82.5, max 141). Overall quality was first assessed using fastqc and adapter contamination was trimmed from the reads using Trimmomatic v.0.39. Trimmed reads were then aligned to the mouse genome assembly (mm10) using the TopHat2 aligner v.2.1.1 based on the gencode.vM2.annotation.gtf. Greater than 80% of all reads were uniquely aligned from each sample (mean 81%, min 77.1%, max 86.6%). Lack of 3’ bias was confirmed with RNA-SeQC. Gene counts for each sample were extracted with featureCounts with an average of 91.2%+/-0.4% of uniquely aligned reads being assigned to a gene. Differential gene analysis was performed in R v3.4.3 using DESeq2 v.1.22.0. Resulting Wald test p-values were adjusted (p-adjust; p-adj) using Benjamini-Hochberg approach for all downstream analysis. Significantly differentially expressed genes (p-adjust <0.05, n=3436, 1730 down, 1706 up) were assessed for functional enrichments using clusterProfiler v3.9.1 or metascape.org. Gene ontology and KEGG pathway enrichments with q-values < 0.05 were considered significant.

### RNAseq analysis of Kcnn1 overexpression

Spinal cords from four B6SJL mice and four Kcnn1-6 transgenic mice were dissected, embedded, and sectioned as above. RNA samples (RIN >7) were recovered as above, and libraries were prepared as described from ∼15 ng of each RNA, except using the NEBNext Directional RNA library kit. Libraries were diluted to 4 nM, pooled and denatured according to the instructions for Illumina NextSeq 1000, and sequenced with settings for single-index, paired-end sequencing with 50 cycles per end. An average of 67 million paired-reads were obtained from each sample (min 59.1, max 75.1). Overall quality was first assessed using fastqc and adapter contamination was trimmed from the reads using Trimmomatic v.0.39. Trimmed reads were then aligned to the mouse genome assembly (mm10) using the STAR aligner v.2.7.11a based on the gencode.vM2.annotation.gtf. Greater than 80% of all reads were uniquely aligned from each sample (mean 85.5%, min 81.3%, max 86.8%). Lack of 3’ bias was confirmed with RNA-SeQC. Gene counts for each sample were extracted with featureCounts with an average of 94.4%+/-0.6% of uniquely aligned reads being assigned to a gene. Differential gene analysis was performed in R v4.4.1 using DESeq2 v.1.44. Resulting Wald test p-values were adjusted (p-adjust; p- adj) using Benjamini-Hochberg approach for all downstream analysis. Significantly differentially expressed genes (p-adjust <0.05, n=1788, 968 down, 820 up) were assessed for functional enrichments using clusterProfiler v4.12.0 or metascape.org. Gene ontology and KEGG pathway enrichments with q-values < 0.05 were considered significant.

## References

Adelman, J.P., Maylie, J., and Sah, P. (2012) Small-conductance Ca^2+^-activated K^+^ channels: Form and function. Annu. Rev. Physiol 74, 245–269.

Anderson, N.S. and Haynes, C.M. (2020) Folding the mitochondrial UPR into the integrated stress response. Trends in Cell Biology 30, 428–439.

Bandyopadhyay, U., Cotney, J., Nagy, M., Oh, S., Leng, J., Mahajan, M., Mane, S., Fenton, W.A., Noonan, J., and Horwich, A.L. (2013) RNA-seq profile of spinal cord motor neurons from a presymptomatic SOD1 ALS mouse. PLoS ONE http://dx.plos.org/10.1371/journal.pone.0053575

Benton, D.C.H., Monaghan, A.S., Hosseini, R., Bahia, P.K., Haylett, D.G. and Moss G.W.J. Small conductance Ca^+2^-activated K^+^ channels formed by the expression of rat SK1 and SK2 genes in HEK 293 cells. (2003) J.Physiol. 533.1, 13–19.

Bommiasamy, H, Back, S.H., Fagone, P., Lee, K., Meshinchi, S., Vink, E., Sriburi, R., Frank, M., Jackowski, S., Kaufman, R.J., and Brewer, J.W. (2009) ATF6alpha induces XBP1-independent expansion of the endoplasmic reticulum. J.Cell Sci.122, 1626–1636.

Caroni, P. (1997) Overexpression of growth-associated proteins in the neurons of adult transgenic mice. J.Neurosci. Meth. 71, 3–9.

Chandra, S., Gallardo, G., Fernández-Chacón, R., Schlüter, O.M., and Südhof, T.C. (2005) α-Synuclein cooperates with CSPα in preventing neurodegeneration. Cell 123, 383–396.

Chiduza, G.N., Garza-Garcia, A., Almacellas, E., De Tito, S., Pye, V.E., van Vliet, A.R., Cherepanov, P., and Tooze, S.A. (2023) ATG9B is a tissue-specific homotrimeric lipid scramblase that can compensate for ATG9A. Autophagy 20, 557–576.

Costa-Mattioli, M. and Walter, P. (2020) The integrated stress response: From mechanism to disease. Science 368, eaat5314

Eura, Y., Miyata, T., and Kokame, K. (2020) Derlin-3 is required for changes in ERAD complex formation under ER stress. Int.J.Mol.Sci. 21, 6146; doi:10.3390/ijms21176146

Ghanem, S.S., Majbour, N.K., Vaikath, N.N, Ardah, M.T., Erskine, D., Jensen, N.M., Fayyad, M., Sudhakaran, I.P, Vasili, E., Melachroinou, K., Abdi, I.Y., Poggiolini, I., Santos, P., Dorn, A., Carloni, P., Vekrellis, K., Attems, J., McKeith, I., Outeiro, T.F., Jensen, P.H., and El-Agnaf, O.M.A. (2022) alpha-synuclein phosphorylation at serine 129 occurs after initial protein deposition and inhibits seeded fibril formation and toxicity. Proc. Natl. Acad. Sci. USA 119, e2109617119.

Gray, J.T., and Zolotukhin, S. (2011) Design and construction of functional AAV vectors. Methods Mol. Biol. 807, 25–46.

Gurney, M.E., Pu, H., Chiu, A.Y., Dal Canto, M.C., Polchow, C.Y., Alexander, D.D., Caliendo, J., Hentati, A., Kwon, Y.W., Deng, H-X., Chen, W., Zhai, P., Sufit, R.L., and Siddique, T. Motor neuron degeneration in mice that express a human Cu,Zn superoxide dismutase mutation. Science 264, 1772–1775.

Hadzipasic, M., Tahvildari, B., Nagy, M., Vian, M., Horwich, A.L., and McCormick, D.A. (2014) Selective degeneration of a physiological subtype of spinal motor neuron in mice with SOD1-linked ALS. Proc. Natl. Acad. Sci. USA 111, 16883–16888.

Hegde, R.S. and Keenan, R.J. (2024) A unifying model for membrane protein biogenesis. *Nature Struct and Molec*. Biol. 31, 1009–1017.

Karampetsou, M., Ardah, M.T., Semitekolou, M., Polissidis, A., Samiotaki, M., Kalomoiri, M., Majbour, N., Xanthou, G., El-Agnaf, O.M.A. and Vekrellis, K. (2017) Phosphorylated exogenous alpha-synuclein fibrils exacerbate pathology and induce neuronal dysfunction in mice. Scientific Reports 7:16533 DOI:10.1038/s41598-017-15813-8

Kim, J.-Y., Grunke, S.D., Levites, Y., Golde, T.E., and Jankowsky, J.L. (2014) Intracerebroventricular viral injection of the neonatal mouse brain for persistent and widespread neuronal transduction. JoVE 91: 51863.

Kohler, M., Hirschberg, B., Bond, C.T., Kinzie, J.M., Marrion, N.V., Maylie, J., and Adelman, J.P. Small-conductance, calcium-activated potassium channels from mammalian brain. Science 273, 1709–1714.

Lee, C.-H. and MacKinnon, R. (2018) Activation mechanism of a human SK-calmodulin channel complex elucidated by cryo-EM structures. Science 360, 508–513.

Long, J.M. and Holtzman, D.M. (2019) Alzheimer Disease: An update on pathobiology and treatment strategies. Cell 179, 312–339.

Martin, L.J., Semenkow, S., Hanaford, A., and Wong, M. (2014) The mitochondrial permeability transition pore regulates Parkinson’s disease development in mutant alpha-synuclein transgenic mice. Neurobiology of Aging 35, 1132–1152.

Melber A. and Haynes, C.M. (2018) UPR^mt^ regulation and output: a stress response mediated by mitochondrial-nuclear communication. Cell Res. 28, 281–295.

Papazian, D.M. (1999) Potassium channels: Some assembly required. Neuron 23, 7–10.

Riley, L.G., Nafisinia, M.,, Menezes, M.J., Nambiar, R., Williams, A., Barnes, E.H., Selvanathan, A., Lichkus, K., Bratkovic, D., Yaplito-Lee, J., Bhattacharya, K., Ellaway, C., Kava, M, Balasubramaniam, S., and Christodoulou, J. (2022) FGF21 outperforms GDF15 as a diagnostic marker of mitochondrial disease in children. Mol.Genetics and Metabolism 135, 63–71.

Schuck, S., Prinz, W.A., Thorn, K.S., Voss, C., and Walter, P. (2009) Membrane expansion alleviatesendoplasmic reticulum stress independently of the unfolded protein response. J. Cell Biol. 187, 525–536.

Shen, Y., Meunier, L., and Hendershot, L.M. (2002) Identification and characterization of a novel endoplasmic reticulum (ER) DnaJ homologue, which stimulates ATPase activity of BiP in vitro and is induced by ER stress*. J.Biol*. Chem. 277, 15947–15956.

Sierksma, A., Escott-Price, V., and De Strooper, B. (2020) Translating genetic risk of Alzheimer’s disease into mechanistic insight and drug targets. Science 370, 61–66.

Singh, S., Brocker, C., Koppaka, V., Ying, C., Jackson, B., Matsumoto, A., Thompson, D.C., and Vasiliou, V. (2013) Aldehyde dehydrogenases in cellular responses to oxidative/electrophilic stress. Free Radic. Biol. Med. 56, 89–101

Stocker, M. (2004) Ca^+2^-activated K^+^ channels: Molecular determinants and function of the SK family. Nature Reviews Neuroscience 5, 758–770.

Tanner, C.M., and Ostrem J.L. (2024) Parkinson’s Disease. NEJM 391, 442–452.

Thomas, E.V., Fenton, W.A., McGrath, J., and Horwich, A.L. (2017) Transfer of pathogenic and non-pathogenic cytosolic proteins between spinal cord motor neurons in vivo in chimeric mice. Proc. Natl. Acad. Sci. USA 114(15):E31329–E3148.

Thomas, E.V., Nagy, M., Fenton, W.A., and Horwich, A.L. (2018) Motor nuclei innervating eye muscles spared in mouse model of SOD1-linked ALS. BioRxiv doi.org/10.1101/304857.

Travers, K.J., Patil, C.K., Wodicka, L., Lockhart, D.J., Weissman, J.S., and Walter, P. (2000) Functional and genomic analyses reveal and essential coordination between the unfolded protein response and ER-associated degradation. Cell 101, 249–258.

Walter, P. and Ron, D. (2011) The unfolded protein response: from stress pathway to homeostatic regulation. Science 334, 1081–1085.

Wang, J., Farr, G.W. Zeiss, C.J., Rodriguez-Gil, D.J., Wilson, J.H., Furtak, K., Rutkowski, D.T., Kaufman, R.J., Ruse, C.I., Yates, J.R. III, Perrin, S., Feany, M.B., and Horwich, A.L. (2009) Progressive aggregation despite chaperone associations of a mutant SOD1-YFP in transgenic mice with ALS. Proc. Natl. Acad. Sci. USA 106, 1392–1397.

Yost, C.S. (1999) Potassium channels. Anesthesiology 90, 1186–1203.

