## Supplemental figure 1 for "Neuronal overexpression of potassium channel subunit Kcnn1 prolongs survival of SOD1-linked ALS and A53T alpha-synuclein mouse models"

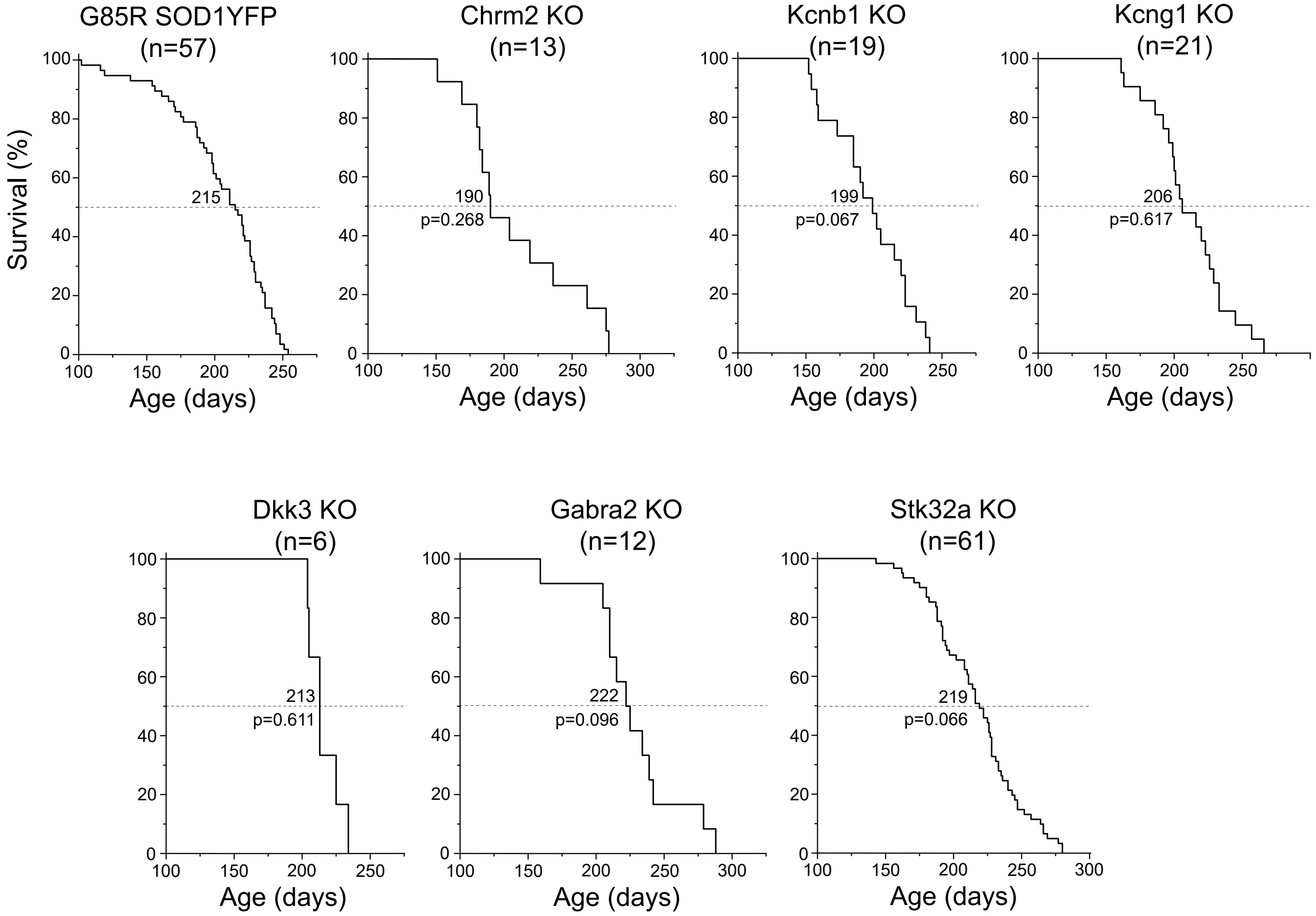


**Supp. Fig. 1.** Kaplan-Meier survival plots for control G85R SOD1YFP homozygous mice alone and homozygously-disrupted of candidate knockout genes: Chrm2 KO, Kcnb1 KO, Kcng1 KO, Dkk3 KO, Gabra2 KO, and Stk32a KO. In normal mice, these six genes exhibited less RNA expression in 3N (spared in ALS) versus 12N and spinal cord motor neurons (susceptible in ALS). Median survival time to paralysis (days) of the candidate knockout strain was compared to the control, and p-values for the plots of the candidates relative to the plot of the control are presented for each candidate.
