## Supplemental figure 2 for "Neuronal overexpression of potassium channel subunit Kcnn1 prolongs survival of SOD1-linked ALS and A53T alpha-synuclein mouse models"

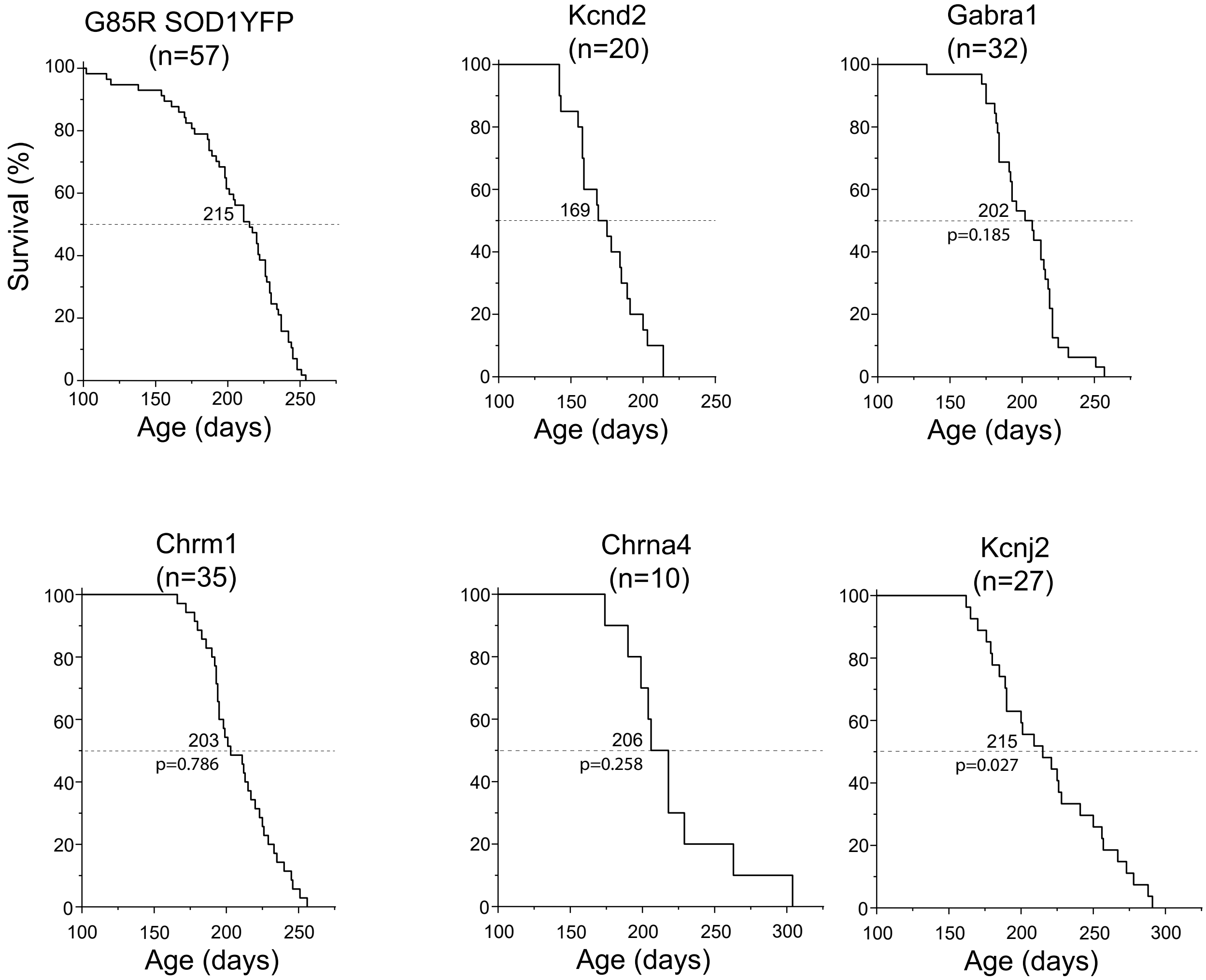


**Supp. Fig. 2.** Kaplan-Meier survival plots for control G85R SOD1YFP homozygous mice alone and hemizygous for transgenic candidate genes, Thy1.2 promoter-driven cDNAs for mouse Kcnd2, Gabra1, Chrm1, Chrna4, and Kcnj2. The five corresponding genes exhibited relatively higher RNA expression in 3N (spared in ALS) vs 12N and spinal cord motor neurons (susceptible in ALS). Median survival to the time of paralysis of the candidate-expressing mice was compared to the control, and p-values for the plots of the candidates relative to the plot of the control are presented for each candidate.
