## Supplemental figure 3 for "Neuronal overexpression of potassium channel subunit Kcnn1 prolongs survival of SOD1-linked ALS and A53T alpha-synuclein mouse models"

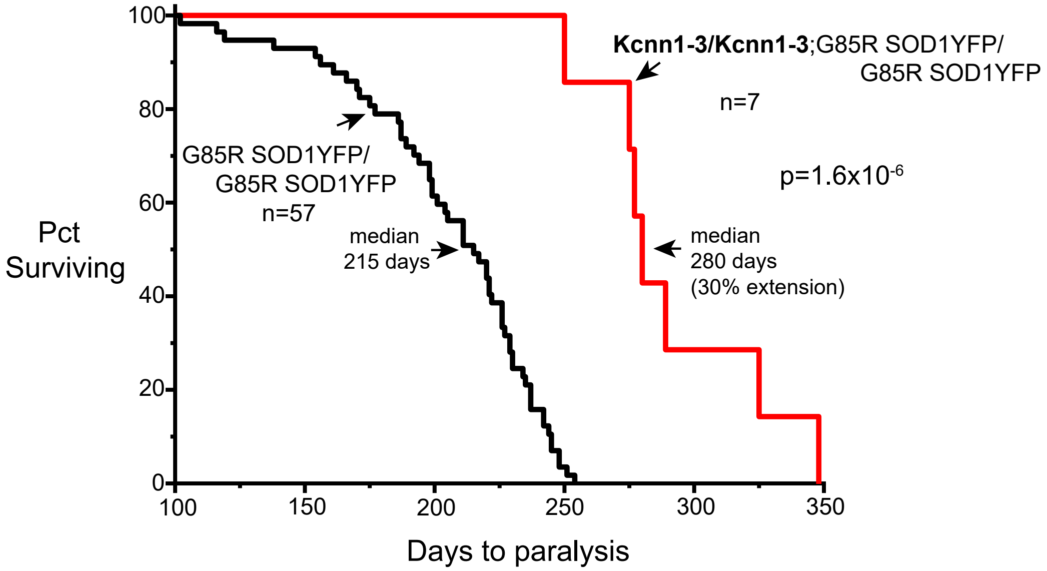


**Supp. Fig. 3.** Kaplan-Meier survival plots for double homozygous Kcnn1-3/Kcnn1-3;G85R SOD1YFP/ G85R SOD1YFP mice, showing that median survival time to paralysis is negligibly further extended as compared with Kcnn1-3/+;G85R SOD1YFP/G85R SOD1YFP (cf Fig.1A) The cohort of homozygous G85R SOD1YFP here is the same as plotted in Fig.1A, with copy number 260-330.
