## Supplemental figure 4 for "Neuronal overexpression of potassium channel subunit Kcnn1 prolongs survival of SOD1-linked ALS and A53T alpha-synuclein mouse models"

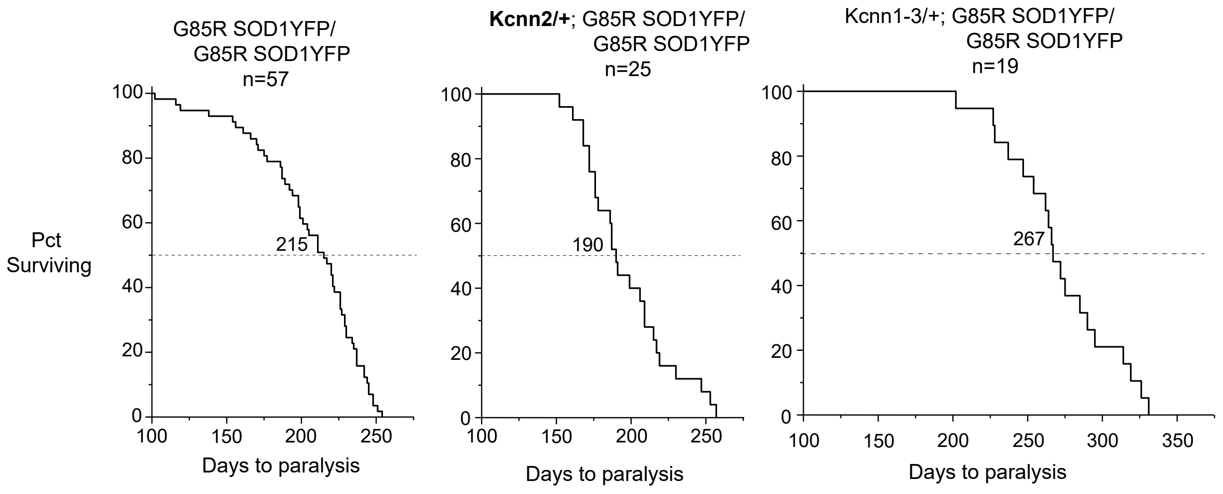


**Supp. Fig. 4.** Transgenic Thy1.2-Kcnn2/+ does not confer extension of survival time to paralysis of G85R SOD1YFP homozygous mice (central Kaplan-Meier survival plot), median survival of 190 days. This plot is compared with cohort plots taken from Fig.1A of homozygous G85R SOD1YFP (left) and Kcnn1-3/+;G85R SOD1YFP/G85R SOD1YFP (right).
