## Supplemental figure 5 for "Neuronal overexpression of potassium channel subunit Kcnn1 prolongs survival of SOD1-linked ALS and A53T alpha-synuclein mouse models"

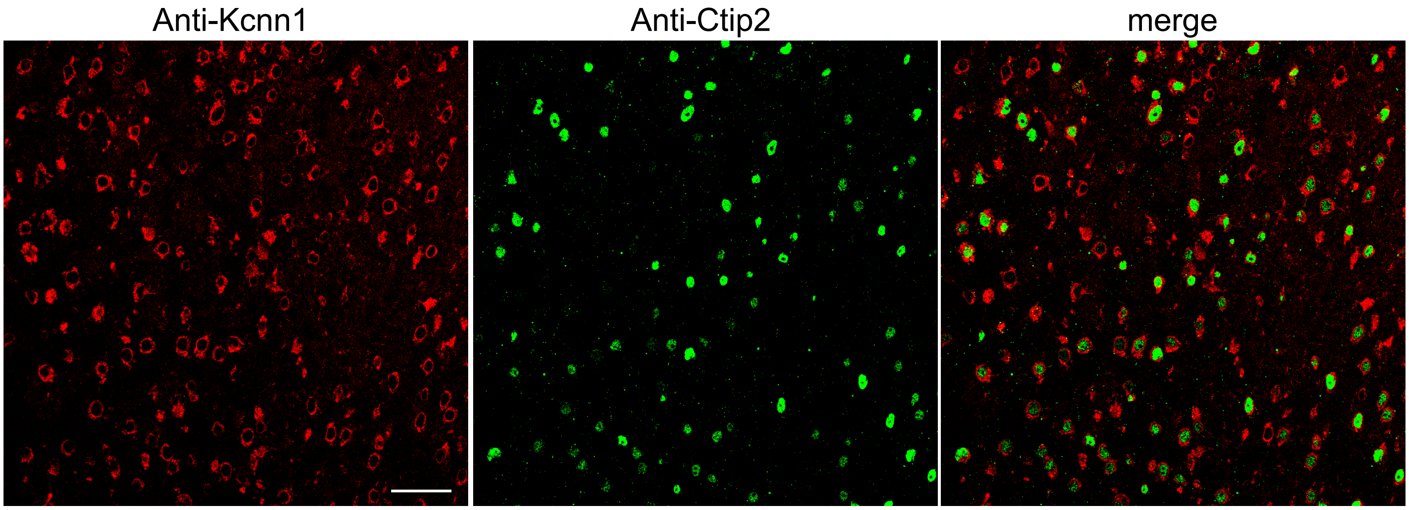


**Supp. Fig. 5.** Cortical motor neurons in layer V of a Kcnn1-3/Kcnn1-3 mouse express Kcnn1-encoded protein. Left panel shows strong cytoplasmic immunostaining of Kcnn1 (as in Fig.3); middle panel shows nuclear staining specific to layer V with anti-Ctip2 antibody; and right panel shows a merge of the two images, demonstrating that at least 50% of the anti-Kcnn1-stained neurons are also Ctip2-reactive.
