## Supplemental figure 6 for "Neuronal overexpression of potassium channel subunit Kcnn1 prolongs survival of SOD1-linked ALS and A53T alpha-synuclein mouse models"

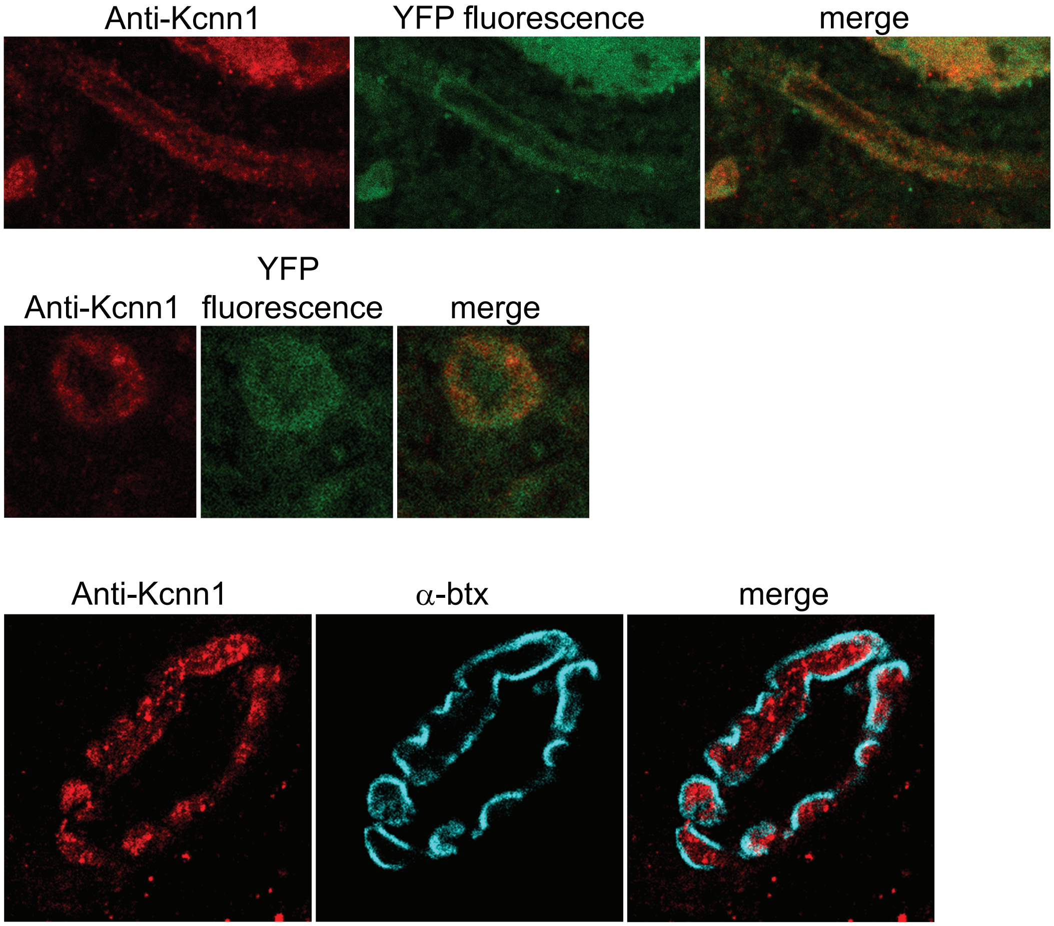


**Supp. Fig. 6.** Kcnn1 localizes also in axons and the presynaptic zone of synapses in a double homozygous Kcnn1-3; G85R SOD1YFP mouse. Top panels, longitudinally sliced axon in cervical spinal cord showing punctate Kcnn1 pattern in exposed edges as well as G85R SOD1YFP, with a merge showing a degree of overlap.
Middle panels, axon in cross-section showing G85R SOD1YFP throughout, including axoplasm, but in Kcnn1 more marginated, in axolemma or myelin sheath.
Lower panels, lumbrical muscle NMJ showing presence of Kcnn1 in presynaptic zone as distinct from bungarotoxin (  btx)-stained postsynaptic zone. The Kcnn1 staining pattern in the presynaptic zone was patchy, resembling the cytoplasmic pattern observed in the various Kcnn1-transgenic spinal cord motor neurons.
