## Supplemental figure 7 for "Neuronal overexpression of potassium channel subunit Kcnn1 prolongs survival of SOD1-linked ALS and A53T alpha-synuclein mouse models"

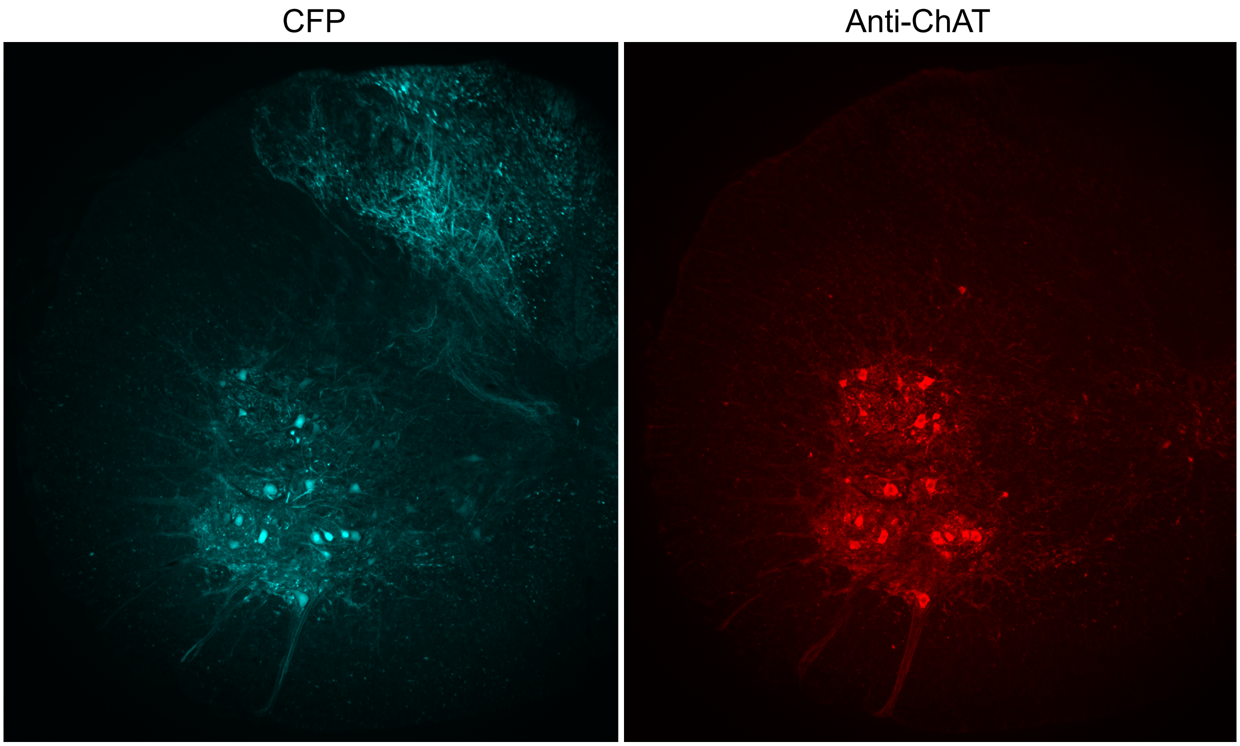


**Supp. Fig. 7.** Cervical spinal cord section of 5 month old B6SJL mouse that had been injected at p0 with scrAAV9CFP virus at 1.2 X 10E13 vp/ml into the lateral ventricles. A 20 micron hemisection is shown with CFP fluorescence in ventral horn neurons (left) and ChAT-positive neurons in the same section identified by immunostaining (right). Approximately 50% of ChAT-positive neurons were positive for CFP.
