## Supplemental figure 8 for "Neuronal overexpression of potassium channel subunit Kcnn1 prolongs survival of SOD1-linked ALS and A53T alpha-synuclein mouse models"

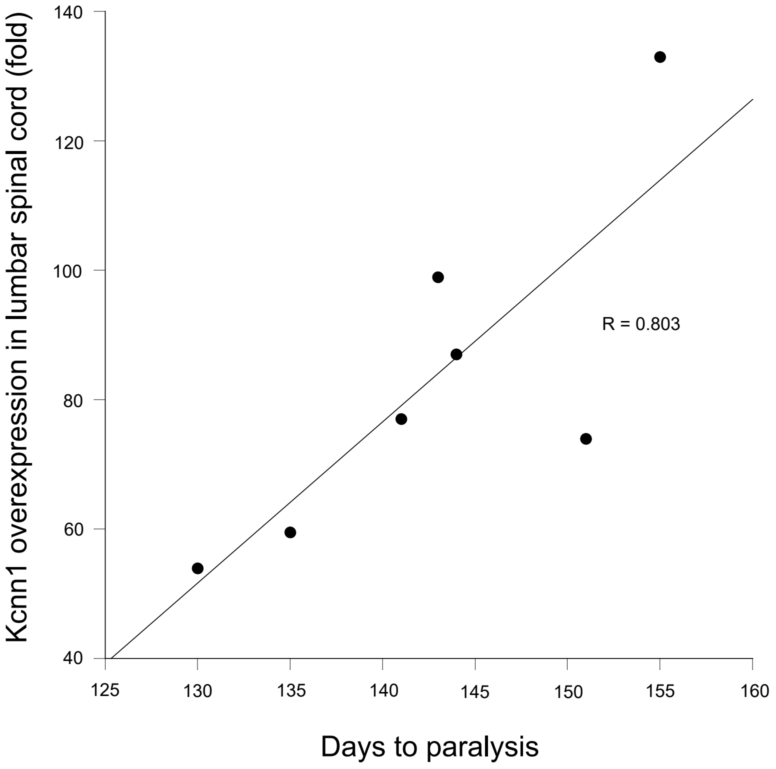


**Supp. Fig. 8.** Correlation of Kcnn1 level of expression measured by qRTPCR with survival time to paralysis. 3 mm lengths of lumbar spinal cord were taken from paralyzing G93A/+ mice that had been injected ICV at P0 with AAV9-CMV-Kcnn1 virus, and RNA prepared for qRTPCR measurement relative to GAPDH. The longest-surviving mouse (155 days) exhibited the greatest fold of overexpression (130X), measured as Kcnn1 RNA level/GAPDH RNA of the injected mouse vs ratio from 3 mm snippet from uninjected B6SJL mouse. The level of overexpression in this mouse and time of extended survival approach those of the median observed for genetic transduction (see Fig. 2B).
