## Supplemental figure 9 for "Neuronal overexpression of potassium channel subunit Kcnn1 prolongs survival of SOD1-linked ALS and A53T alpha-synuclein mouse models"

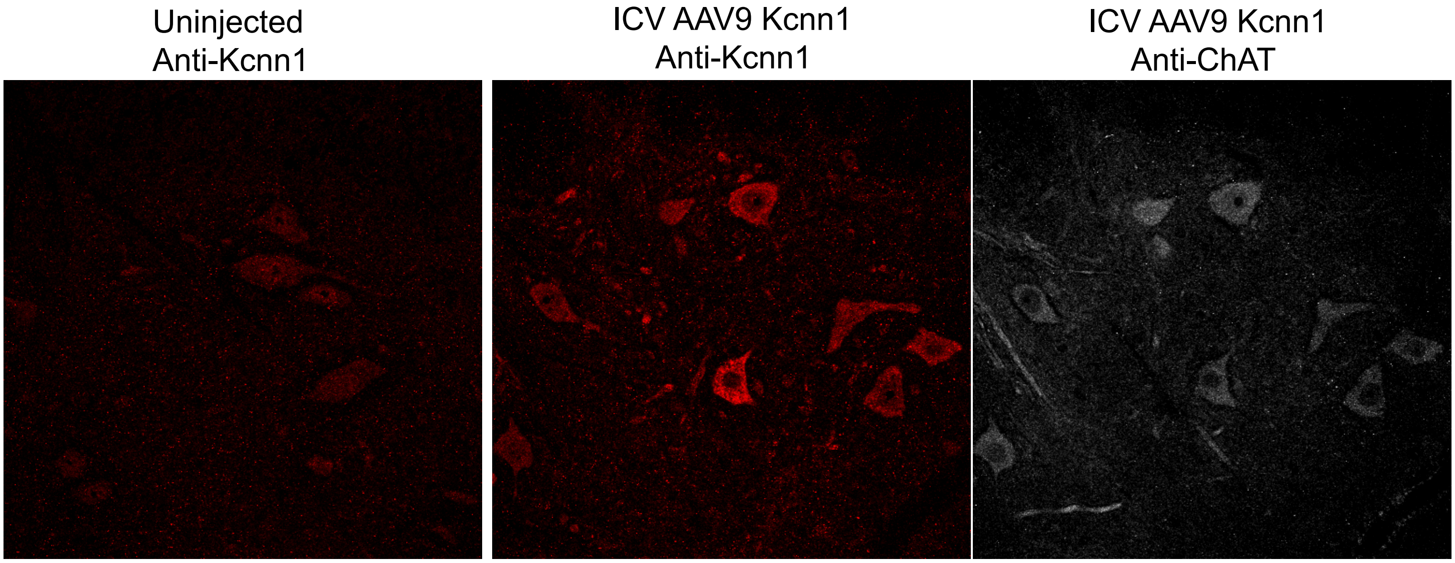


**Supp. Fig. 9.** Anti-Kcnn1 immunostaining of transduced motor neurons in cervical spinal cord section from 155 day old ICV-injected mouse of SF8 (middle panel), showing strong immunostaining relative to uninjected B6SJL mouse (left panel). Most motor neurons in the section from this mouse were Kcnn1-transduced as shown by comparison with anti-ChAT-staining (right panel).
