## Supplemental figure 10 for "Neuronal overexpression of potassium channel subunit Kcnn1 prolongs survival of SOD1-linked ALS and A53T alpha-synuclein mouse models"

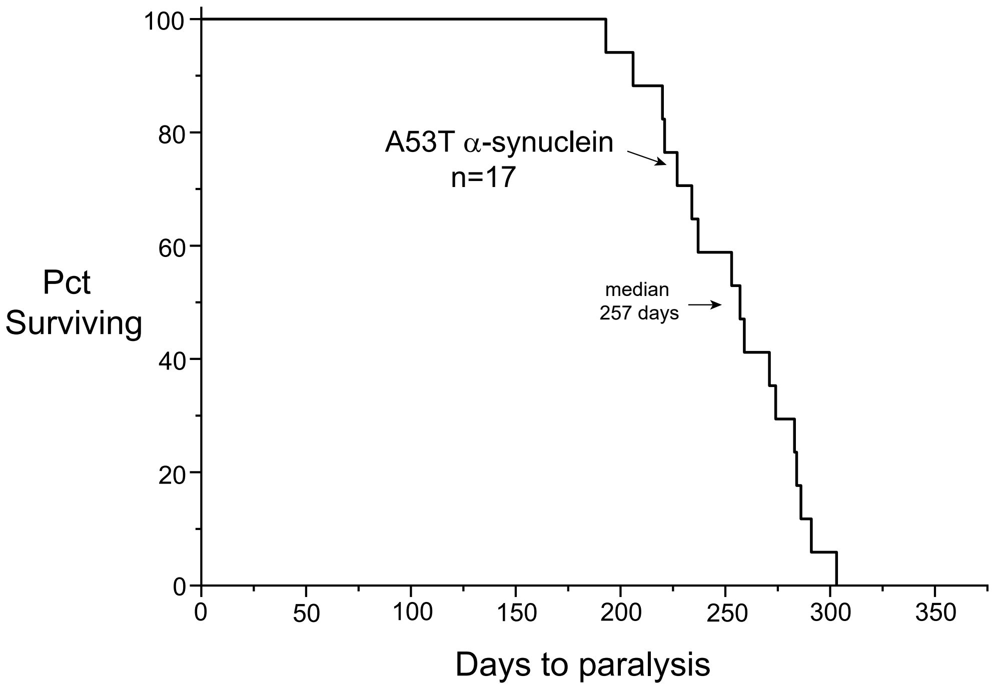


**Supp. Fig. 10.** Kaplan-Meier survival plot showing time of survival to endstage motor compromise of Thy1.2-A53T alpha-synuclein transgenic mice (B6SJL background). Median survival was 257 days.
