## Supplemental figure 11 for "Neuronal overexpression of potassium channel subunit Kcnn1 prolongs survival of SOD1-linked ALS and A53T alpha-synuclein mouse models"

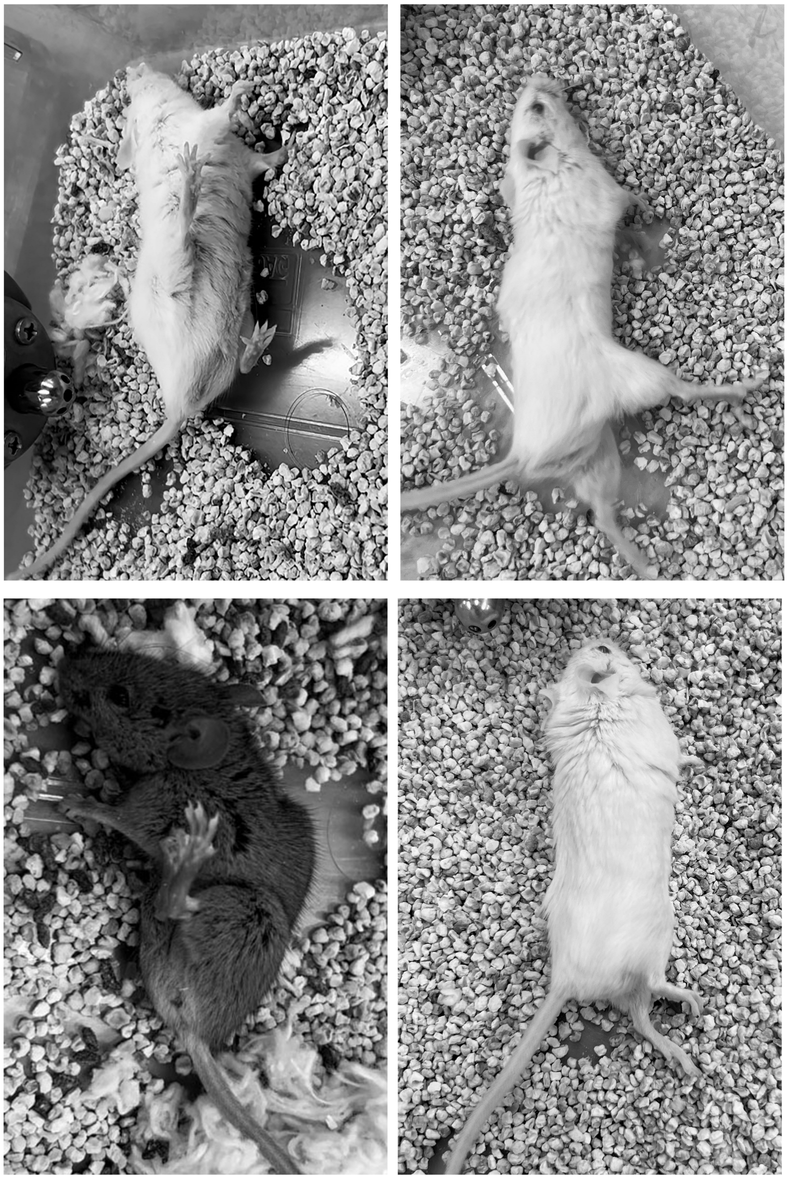

**Supp. Fig. 11.** Immobility and inability to maintain upright position at endstage of A53T alpha-synuclein/B6SJL transgenic mice. Note variability of position of extremities (with lower extremity spasticity manifest in mouse at lower left).
