## Supplemental figure 12 for "Neuronal overexpression of potassium channel subunit Kcnn1 prolongs survival of SOD1-linked ALS and A53T alpha-synuclein mouse models"

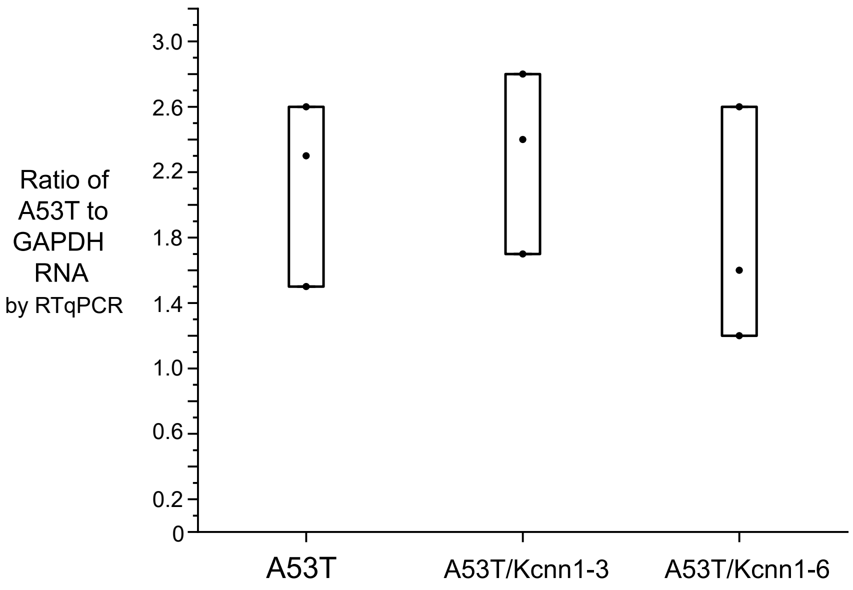


**Supp. Fig. 12.** A53T alpha-synuclein RNA levels in brain are not substantially inhibited by presence of Kcnn1-3 or Kcnn1-6 transgenes. Total RNA was prepared from one side of the brain of A53T/+ B6SJL or the respective double transgenic Kcnn1/A53T mice and level of alpha-synuclein RNA relative to GAPDH determined by qRTPCR. There was considerable variability in A53T level among the 3 mice of each genotype, but no overall diminution of alpha synuclein RNA was apparent.
